# Macrocyclization of an all-D linear peptide improves target affinity and imparts cellular activity: A novel stapled α-helical peptide modality

**DOI:** 10.1101/767673

**Authors:** Srinivasaraghavan Kannan, Pietro G. A. Aronica, Simon Ng, Dawn Thean, Yuri Frosi, Sharon Chee, Jiang Shimin, Tsz Ying Yuen, Ahmad Sadruddin, Hung Yi Kristal Kaan, Arun Chandramohan, Jin Huei Wong, Yaw Sing Tan, Fernando J. Ferrer, Prakash Arumugam, Yi Han, Shiying Chen, Christopher J. Brown, Charles W. Johannes, Brian Henry, David P. Lane, Tomi K. Sawyer, Chandra S. Verma, Anthony W. Partridge

**Author notes:** Corresponding authors: Srinivasaraghavan Kannan, Bioinformatics Institute (A*STAR), 30, Biopolis Street, 07-01 Matrix, Singapore 138671, Singapore. Chandra S. Verma, Bioinformatics Institute (A*STAR), 30, Biopolis Street, 07-01 Matrix, Singapore 138671, Singapore. Anthony Partridge, MSD International, Translation Medicine Research Centre, 8 Biomedical Grove, #04-01/05 Neuros Building,Singapore. 138665, Singapore.

## Abstract

Peptide-based inhibitors hold great potential for targeted modulation of intracellular protein-protein interactions (PPIs) by leveraging vast chemical space relative to primary structure *via* sequence diversity as well as conformationally through varying secondary and tertiary structures. However, the development of peptide therapeutics has been hindered because of their limited conformational stability, proteolytic sensitivity and cell permeability. Several contemporary peptide design strategies address these issues to varying degrees. Strategic macrocyclization through optimally placed chemical braces such as olefinic hydrocarbon crosslinks, commonly referred to as staples, may address these issues by i) restricting conformational freedom to improve target affinities, ii) improving proteolytic resistance, and iii) enhancing cell permeability. Conversely, molecules constructed entirely from D-amino acids are hyper-resistant to proteolytic cleavage, but generally lack conformational stability and membrane permeability. Since neither approach is a complete solution, we have combined these strategies to identify the first examples of all-D α-helical stapled and stitched peptides. As a template, we used a recently reported all D-linear peptide that is a potent inhibitor of the p53-Mdm2 interaction, but is devoid of cellular activity. To design both stapled and stitched all-D-peptide analogues, we used computational modelling to predict optimal staple placement. The resultant novel macrocyclic all D-peptide was determined to exhibit increased α-helicity, improved target binding, complete proteolytic stability and, most notably, cellular activity.

## Introduction

Protein-protein interactions (PPIs) are central to most biological processes and are often dysregulated in disease [1, 2]. Therefore, PPIs are attractive therapeutic targets for novel drug discovery. However, in contrast to the deep protein cavities that typically accommodate small molecules, PPI surfaces are generally large and flat, and this has contributed to the limited success in developing small molecule inhibitors for PPI targets [3]. The realization that ~40% of all PPIs are mediated by relatively short peptide motifs gave rise to the possibility of developing peptide-based inhibitors that would compete orthosterically for the interface between ligand–target cognate partners [4]. When taken out of the protein ligand context and synthesized, such peptides may often be unstructured and intrinsically disordered, yet able to achieve their biologically-relevant conformation upon protein target binding [4]. However, for intracellular targets, the peptide modality may be challenging due to proteolytic sensitivity, low conformational stability (yielding weak affinities and off-target effects), and poor cell permeability (further limiting engagement of intracellular targets and/or oral bioavailability) [5–11]. To address these issues, several strategies have been pursued, including macrocyclization and modifications of the peptide backbone to yield molecules with improved biological activities and pharmacokinetic properties as well as constraining the peptide into its biologically-relevant conformation that binds its target [5–13]. First, by constraining the peptides toward their bound conformations, entropic penalties upon binding are reduced, thus improving binding constants as well as potentially decreasing the opportunity for unwanted off-target effects. Secondly, macrocyclization may confer varying degrees of proteolytic resistance by modifying key backbone and/or side-chain structural moieties in the peptide. Thirdly, macrocyclization may enhance cell permeability, such as through increased stability of intramolecular hydrogen bonding to reduce the desolvation penalty otherwise incurred in the transport of peptides across an apolar cell membrane. Amongst the several cyclization techniques described, stapling *via* olefin metathesis using a non-proteogenic amino acid such as alpha-methyl alkenyl side chains has proven to be very effective [13–18], particularly when the desired secondary structure of the peptide macrocycle is helical. Stapling requires incorporation of the appropriate unnatural amino acid precursors to be placed at appropriate locations along the peptide sequence such that they do not interfere with the binding face of the helix. The linkers can be of different types, and can span different lengths, resulting in *i,i+3 i,i+4, i,i+7* staples. Although they have largely been used to stabilize helical conformations, recent studies have also applied RCM strategies to non-helical peptides [19, 20].

The stapled peptide strategy has been successfully applied to inhibit several PPIs of therapeutic potential including, BCL-2/Mcl-1 family [21–24], β-catenin–TCF [25], Rab–GTPase-Effector [26], ERα–coactivator protein [27], Cullin3–BTB [28], VDR–coactivator protein [29], eIf4E [30] and p53–Mdm2/Mdm4 [31–34]. Noteworthy, in the case of p53–Mdm2/Mdm4, a dual selective stapled peptide (ALRN-6924) has been further successfully advanced to phase II clinical trials [35–37]. Although this example is unquestionably encouraging for the advancement of stapled peptides into the clinic, challenges still remain. Amongst these, engineering molecules with sufficient proteolytic stability for sustained target binding and cellular activity is critical. Indeed, although stapling L-amino acid peptides can confer resistance to protease-mediated degradation, the effect is often not complete, and may affect residues located outside of the macrocycle [38–40]. On the other hand, all-D peptides are hyperstable against proteolysis and have been engineered with strong binding affinity against a variety of targets including p53–Mdm2 [41–42], VEGF–VEGF-receptor [43], PD-1–PD-L1 [44], and human immunodeficiency virus type 1 (HIV-1) entry [45]. Unfortunately, although all-D peptides are intrinsically hyperstable to proteolysis, they generally lack membrane permeability and cellular activity.

We hypothesized that combining both strategies, ie all-D-peptide and helical macrocyclization might provide synergistic results. Thus, we embarked on introducing a hydrocarbon staple into an all-D peptide inhibitor of the p53 - Mdm2/Mdm4 interaction. p53 is a key tumour suppressor protein, which primarily functions as a DNA transcription factor, which is commonly inactivated in cancer and normally plays a crucial role in guarding the cell in response to various stress signals through the induction of cell cycle arrest, apoptosis or senescence [46]. Mechanisms that frequently result in the inactivation of p53 and tumorigenesis include increased expression of the p53-negative regulators Mdm2 and Mdm4. Both Mdm2 and Mdm4 attenuate p53 function by interacting directly with p53 and preventing its interaction with the relevant activation factors required for transcription e.g. dTAF_II_, hTAF_II_. In addition, they are both E3 ligase components and target p53 for proteosomal mediated degradation. Mdm4, unlike Mdm2, has no intrinsic E3 ubiquitin ligase activity. Instead Mdm4 forms heterodimeric complexes with Mdm2 whereby it stimulates the ubiquitin activity of Mdm2. As a result, p53 activity and protein levels are acutely suppressed by Mdm2 and Mdm4 overexpression. Development of inhibitors to disrupt the interactions of p53 with either Mdm2 or Mdm4, or both, are therefore highly desirable as they will prevent p53 degradation and restore a p53 dependent transcriptional anti-tumour response [47,48].

The structural interface of the p53 Mdm2/Mdm4 complex is characterized by an α-helix from the N-terminal transactivation domain of p53 which binds into a hydrophobic groove on the surface of the N-terminal domain of both Mdm2 and Mdm4. Three hydrophobic residues, Phe19, Trp23 and Leu26 from p53 are critical determinants of this interaction and project deeply into the Mdm2 interaction groove [Figure 1A]. The isolated p53 peptide is largely disordered, morphing into an α-helical conformation upon binding. There are several examples of small molecules, peptides and biologics that mimic these interactions and compete for Mdm2/4 binding, with the release of p53 [37]. However, a large majority of the small molecules developed exhibit little affinity and activity against Mdm4, which possesses several distinct structural differences in the p53 peptide binding groove compared to Mdm2. Although several Mdm2 specific molecules have entered initial clinical trials, they have largely been met with dose limiting toxicities in patients [37]. Overexpression of Mdm4 in tumours has been demonstrated to attenuate the effectiveness of Mdm2 specific compounds, presumably through the maintenance of heterodimeric complexes of Mdm2 and Mdm4 that inhibit and target p53 for proteosomal degradation. Mdm2-selective inhibitors may also induce higher levels of Mdm4. This highlights the importance of targeting both proteins simultaneously to achieve efficient activation of p53 to achieve an optimal therapeutic response. The ALRN-6924 a dual binding Mdm2/4 L-amino acid stapled peptide is currently in clinical trial and showing promise in terms of tumour response and low toxicity [49].

**Figure 1:**
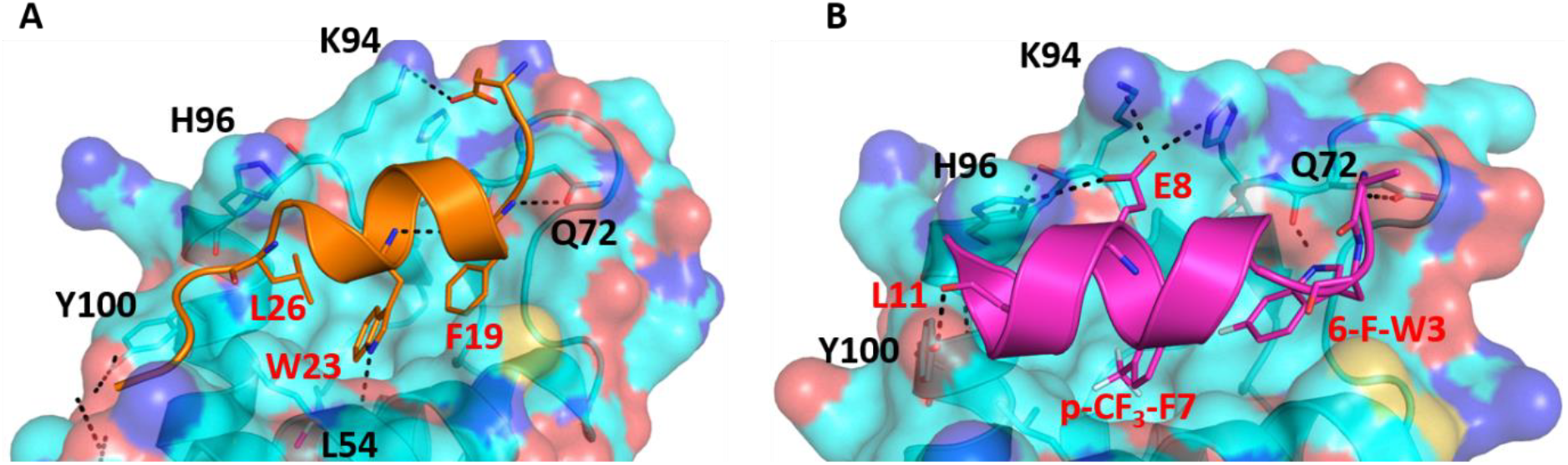
Structural representation of crystal structures of (A) p53-Mdm2 (PDB: 1YCR) and (B) ^d^PMI-δ – Mdm2 (PDB: 3TPX) complex. Mdm2 is shown as surface and bound peptide is shown as cartoon with interacting residues highlighted in sticks. Hydrogen bond interactions are shown as dotted lines (black).

^d^PMI-δ, the all-D linear peptide that served as our starting point, was derived from a mirror image phage display screen reported by Liu et al [41]. Specifically, they reported several 12-mer D-peptide antagonists of Mdm2 (termed ^d^PMI-α, β, γ) that bind with affinities as low as 35 nM and are resistant to proteolytic degradation. ^d^PMI-δ is a corresponding analogue that was modified with two unnatural amino acids (6-F-^d^Trp3 and *p-*CF_3_-^d^Phe7) to improve the Mdm2 binding K_d_ to 220 pM [50]. Crystal structures [50] of the complex between this peptide and the N-terminal domain of Mdm2 showed that the peptide was bound in a conformation similar to that adopted by the wild-type peptide (the all-L amino acid peptide derived from p53). The helix, as expected, was left-handed and projected the side chains of 6-F-^d^Trp3, *p*-CF_3_-dPhe7 and ^d^Leu11 into the hydrophobic pocket of Mdm2 [Figure 1B], in conformations similar to those adopted by the side chains of Phe19, Trp23 and Leu26 in the wild type peptide [Figure 1A]. Unfortunately, this peptide lacked cell permeability, but did activate p53 in cells when delivered using nano-carriers [42]. Given our recent success in the cellular inhibition of Mdm2 and activation of p53 by stapled peptides (containing L-amino acids) [33, 51–53], we questioned whether we could achieve cellular permeation by stapling the D-peptide. We report here the computational design, synthesis and biological evaluation of stapled versions of ^d^PMI-δ. These modifications improved binding and imparted cell permeability to result in disruption of the p53-Mdm2/Mdm4 interaction and ultimately upregulate p53 activity. As an extension of our work, we show that a bicyclic (stitched) version of these peptides demonstrates superior binding and cellular properties relative to the stapled peptide precursors.

## Results

### Conformational landscape of ^d^PMI-δ peptide in apo and Mdm2-bound states

We sought to rationally design stapled ^d^PMI-δ analogues that would stabilize helical structure and preserve or enhance binding affinities. Accordingly, we applied molecular dynamics (MD) simulations to the published co-crystal structure of the Mdm2-^d^PMI-δ complex to understand its structural details critical for the maintenance of the binding motif. During the simulation, the bound conformation of the ^d^PMI-δ peptide remained stable with an RMSD of < 2Å relative to its starting conformation [Figure 2A]. The bound ^d^PMI-δ peptide retained its crystallographic α-helical conformation throughout the simulation (>95% α-helicity). The peptide bound state of Mdm2 also remained stable with an RMSD of < 2Å [Figure 2B]. The bound conformation of the peptide is stabilized by hydrogen bonds and hydrophobic interactions. A hydrogen bond observed in the crystal structure between the side chain N of 6-F-^d^Trp3 and the backbone O of Gln72 [Figure 1B], is preserved in ~80% of the simulation. Other hydrogen bonds seen in the crystal structure and reflected in the simulations but for shorter durations included i) the side chains of Gln72(Mdm2) and ^d^Thr1(^d^PMI-δ), ii) the side chains of Lys94/His96(Mdm2) and ^d^Glu8(^d^PMI-δ), iii) the side chains of His96/Tyr100(Mdm2) and the backbone carbonyl of ^d^Leu11(^d^PMI-δ) [Figure 1B]. As expected, the three critical residues 6-F-^d^Trp3, p-CF3-^d^Phe7 and ^d^Leu11 from ^d^PMI-δ were buried into the hydrophobic binding pocket in Mdm2 [Figure 2C] throughout the simulation. Peptide design was also informed by understanding the conformational landscape of the free ^d^PMI-δ peptide in solution. Simulations were carried out starting from the bound conformation of the peptide extracted from the crystal structure of the Mdm2-^d^PMI-δ complex. Biasing Potential Replica Exchange MD (BP-REMD), a Hamiltonian Replica Exchange Method that has been used successfully to explore peptide landscapes [54, 55], was used to enhance the conformational sampling of the peptide. Unsurprisingly, the free peptide exhibited increased flexibility with RMSD ranging between 2-6 Å [Figure 2D]. The two peaks (3-4 Å and 6 Å) correspond to the partially folded and unfolded states of the peptide, a rapid loss in α-helicity is seen resulting in a state where only ~21% of the sampled conformations are alpha helical. This prediction was experimentally confirmed by circular dichroism (CD) spectroscopy, which showed the peptide was ~20.4% helical in solution [Figure 2E]. This was also expected and consistent with the linear peptides derived from the natural p53 sequence.

**Figure 2:**
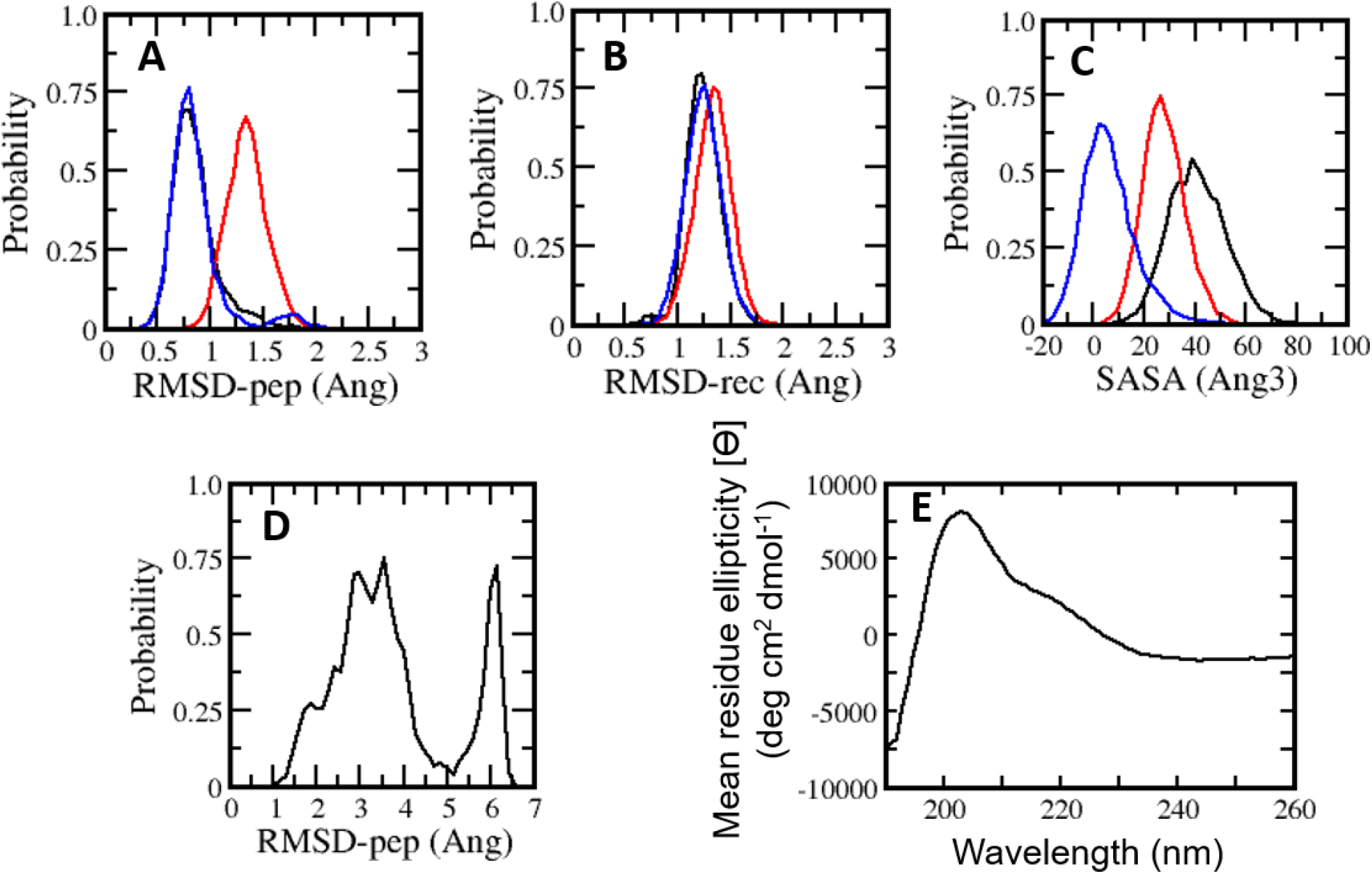
Probability distributions of RMSD (root mean square deviation) (A, B, three colours correspond to triplicate simulations), SASA (solvent accessible surface area) of 6-F-W3 (black), p-CF3-F7 (red), L11 (blue) (C) of conformations sampled during MD simulations of the ^d^PMI-δ – Mdm2 complex. (D) Probability distribution of RMSD of peptide conformations sampled during BP-REMD simulations in the absence of Mdm2. (E) CD spectra of the ^d^PMI-δ peptide Note that this spectra is inverted, as expected for a peptide consisting of D-amino acids only

### Design and Synthesis of stapled D-peptides

Relative to the above simulations, we sought to design stapled analogues of ^d^PMI-δ that would maximize helicity in solution and maintain target binding. To identify appropriate positions on the ^d^PMI-δ peptide for the introduction of the hydrocarbon linkers, we sought to determine residues that, upon mutation, would result in minimal perturbation to the peptide-Mdm2 interaction. The overall binding energy of the peptide to Mdm2 during the MD simulations is decomposed into the energetic contributions of each residue of the peptide. Unsurprisingly, 6-F-^d^Trp3, p-CF3-^d^Phe7 and ^d^Leu11 are the major contributors to the total binding energy followed by ^d^Tyr4 and ^d^Leu10 [Figure 3A]. The contributions from the other 7 residues are either negligible or even slightly destabilizing. We next carried out computational alanine scans of the residues of the peptide by mutating each residue to D-alanine and computing the change in the binding energy for each conformation sampled during the MD simulation and averaging the changes [Figure 3B]. The results mirror the residue-wise contributions [Figure 3A] in that the D-alanine mutations were most deleterious at positions that contributed most, i.e. 6-F-^d^Trp3 and p-CF3-^d^Phe7 of ^d^PMI-δ (> ~10 kcal/mol) [Figure 3B] while substitutions at positions ^d^Tyr4, ^d^Glu8 and ^d^Leu11 resulted in loss of ~2-5 kcal/mol in the overall binding energy [Figure 3B]. In contrast the other positions were quite tolerant to D-Ala substitutions. Overall, these studies suggest 7 positions where staples could be incorporated without significant perturbations to target binding. The incorporation of staples requires careful selection of sidechains with appropriate stereochemistry. As stapling of the left-handed alpha-helices that are formed by all-D peptide has not been conducted previously, we first needed to select the appropriate stereocenters. We reasoned that the stereo-centers should be a mirror-image of the standard strategies that have proven effective for stapling right-handed alpha-helices (*i.e.*, S5 to S5 for (i, i+4) linkages, and R8 to S5 for (i, i+7) linkages). Accordingly, we choose to employ R5 to R5 and S8 to R5 linkages. Using these linkages and the simulations to guide staple placement, we designed several stapled versions of ^d^PMI-δ (details are shown in Figure 4).

**Figure 3:**
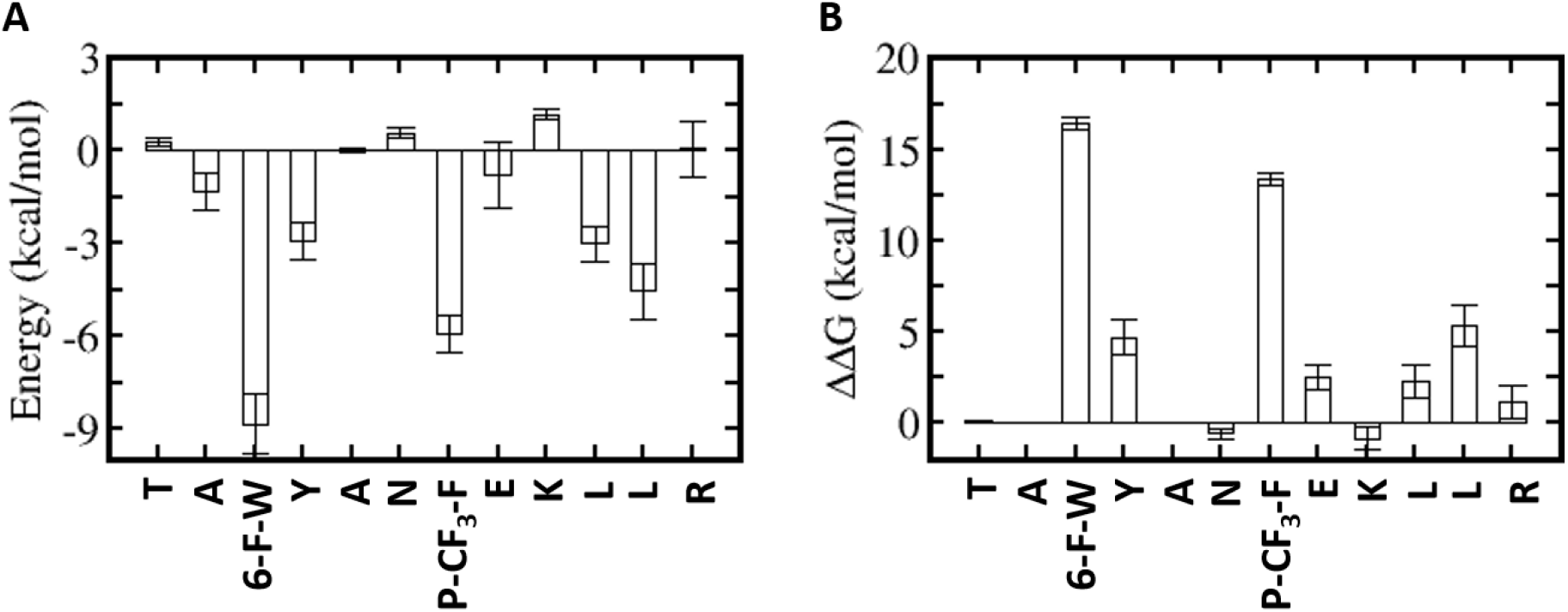
Energetic analysis of the MD simulations of the ^d^PMI-δ – Mdm2 complex. (A) Binding free energy contributions of the ^d^PMI-δ peptide residues. B) Computational alanine (D-ala) scanning of ^d^PMI-δ peptide residues.

**Figure 4:**
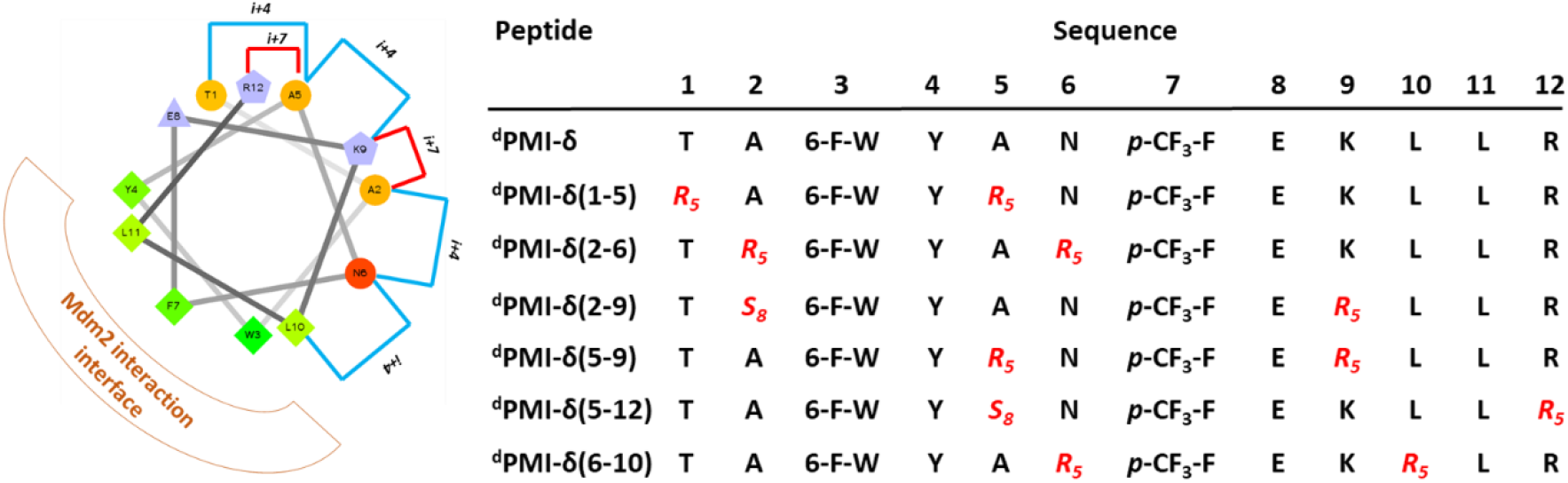
A helical wheel representation of the ^d^PMI-δ template sequence used for the design of stapled peptides. Residues that are linked through all hydrocarbon linkers *i, i+4* and *i, i+7* are highlighted in blue and red respectively. Sequences of the stapled ^d^PMI-δ peptides are shown on the right.

### Peptide stapling increases helicity

BP-REMD simulations suggested that all of the designed stapled ^d^PMI-δ analogues should have increased solution-based helicity. Specifically, we predicted solution helicities between 24-39%; values that were increased compared to the predicted and measured values of ~21% for the unstapled parent sequence (*vida supra*). The values for the stapled analogues agreed well with those obtained experimentally via CD spectroscopy (ranging from 24.5 % to 40%) [Figure 5, Table 1]. This increase in helicity upon stapling mirrors what has been reported for stapling all-L amino acid peptides [14, 56].

**Table 1:** Secondary structure, binding and cellular activity of stapled ^d^PMI-δ peptides determined through various biophysical and biochemical methods.

| Peptide | CD<br>(%helicity) | FP<br>Mdm2<br>Kd (nM) | FP<br>Mdm4<br>Kd (nM) | SPR<br>Kd (nM) | P53 activity<br>EC<sub>50</sub> (μM)<br>(2% FBS,<br>16hrs) | LDH<br>release<br>EC<sub>50</sub> (μM)<br>(2% FBS,<br>16hrs) | Counter<br>screen<br>EC<sub>50</sub> (μM)<br>(2% FBS,<br>16hrs) |
| --- | --- | --- | --- | --- | --- | --- | --- |
| <sup>d</sup>PMI-δ | 20.4 | 36.2 | 43.8 | < 1 | > 50 | > 50 | > 50 |
| <sup>d</sup>PMI-δ(1-5) | 24.7 | 13.2 | 4.5 | < 1 | 13.7 | 24 | > 50 |
| <sup>d</sup>PMI-δ(2-6) | 46.7 | 10000 | 10000 | > 500 | > 50 | 17.7 | 13.9 |
| <sup>d</sup>PMI-δ(2-9) | 26.5 | 3350 | 10000 | 7.4 | > 50 | > 50 | > 50 |
| <sup>d</sup>PMI-δ(5-9) | 38.0 | 10000 | 10000 | 29 | > 50 | 46.5 | 47 |
| <sup>d</sup>PMI-δ(6-10) | 39.9 | 10000 | 10000 | 59 | 30.3 | 18.5 | 39 |
| <sup>d</sup>PMI-δ(5-12) | 31.8 | 38.75 | 29.5 | < 1 | 21.3 | > 50 | > 50 |
| ATSP-7041 | 49.6 | 1.8 | 4.5 | < 1 | 4.2 | > 50 | > 50 |

**Figure 5:**
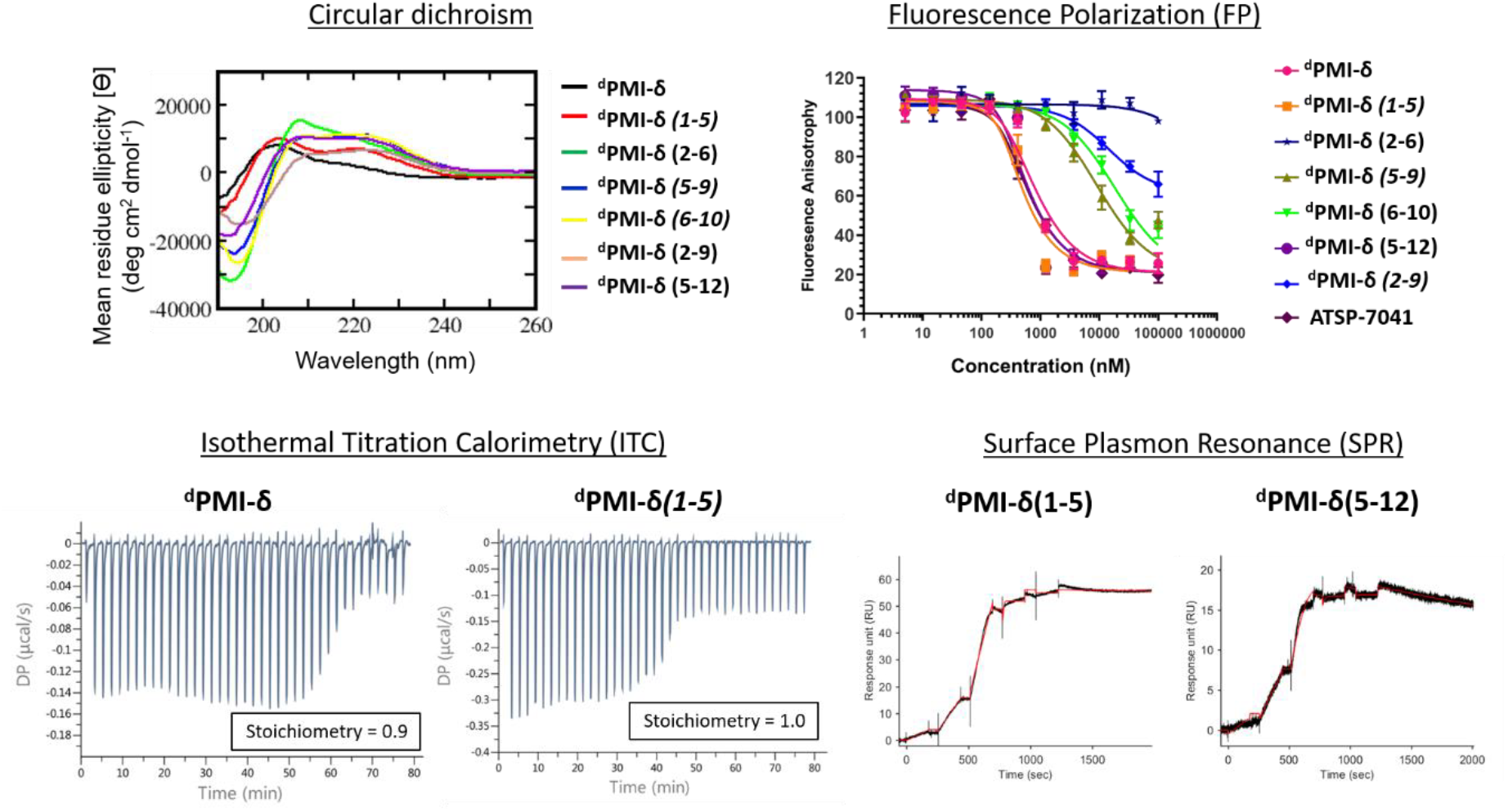
Secondary structure analysis and binding activity of stapled ^d^PMI-δ peptides toward Mdm2 protein determined through various biophysical and biochemical methods. Circular dichroism (CD) spectra of stapled ^d^PMI-δ peptides. Note that this spectra is inverted, as expected for a peptide consisting of D-amino acids only. Fluorescence polarization binding analysis of stapled ^d^PMI-δ peptides and Mdm2 protein. Binding activity of peptides toward Mdm2 protein determined through ITC and binding activity of peptides toward Mdm2 protein determined through SPR.

### Stability and binding affinity are improved upon peptide stapling

We next carried out MD simulations of the stapled ^d^PMI-δ peptides bound to Mdm2. Using the linear ^d^PMI-δ peptide/Mdm2 co-crystal structure as a starting point, staples were modelled into the all-D peptide at 6 sets of residues and subject to MD simulations. The stapled peptides remained stable during the MD simulations and remained largely (~95%) helical. The three critical residues 6-F-^d^Trp3, p-CF3-^d^Phe7 and ^d^Leu11 remained buried in the hydrophobic pocket/binding site of Mdm2 [Figure 6]. The hydrocarbon linkers remained largely exposed to solvent without engaging the Mdm2 surface; this contrasts with some of the L-amino acid stapled peptides where the staples contributed to the binding by engaging with the surface of Mdm2 [33, 52].

**Figure 6:**
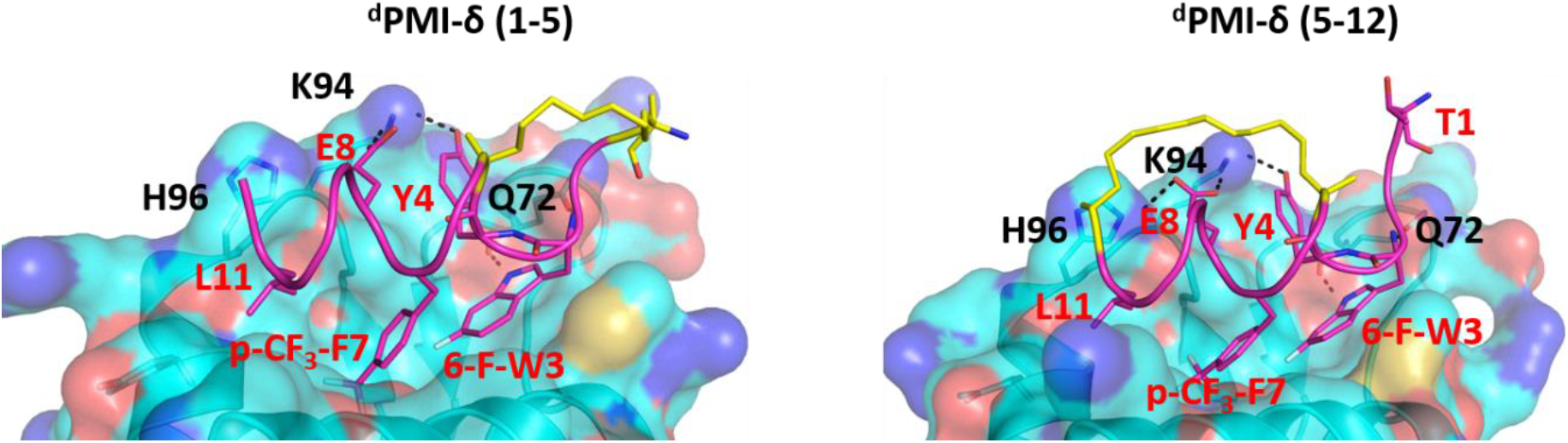
Structural representation of MD snapshot of ^d^PMI-δ(1-5) -Mdm2 (left) and ^d^PMI-δ(5-12)– Mdm2 (right) complex. Mdm2 is shown as surface and bound peptide is shown as cartoon with interacting residues highlighted in sticks. The hydrocarbon linker is highlighted in yellow. Hydrogen bond interactions are shown as dotted lines (black).

Next, the ability of these peptides to bind Mdm2 was measured using fluorescence polarization (FP), surface plasmon resonance (SPR), and isothermal titration calorimetry (ITC). For the binding assays, Mdm2 (residues 17-125) was used in conjunction with linear and stapled ^d^PMI-δ peptides. We used a stapled all-L peptide, ATSP-7041 [34], a validated Mdm2 binder, as a positive control and found that it binds strongly to Mdm2 with Kd of ~2 nM in this version of our FP assay. In our hands, the linear ^d^PMI-δ peptide displayed strong affinity for Mdm2 with Kd of ~36 nM. Two of the six stapled ^d^PMI-δ peptides displayed strong affinity towards Mdm2, two stapled peptides ^d^PMI-δ*(1-5)* and ^d^PMI-δ*(5-12)* binding with Kd of ~13 and ~39 nM respectively. In contrast, peptides ^d^PMI-δ*(2-6)*, ^d^PMI-δ*(5-9)*, ^d^PMI-δ*(6-10)* and ^d^PMI-δ*(2-9)* displayed no binding in the FP assay (Figure 5, Table 1). The SPR binding data mirrored the FP assay, with the linear and stapled ^d^PMI-δ (1-5 staple, and 5-12 staple) binding strongly with Kd of < 1nM, whereas the other stapled ^d^PMI-δ peptides displaying reduced affinity with ^d^PMI-δ (2-6) as a non-binder (Figure 5, Table 1). Models provided an explanation for the lack of binding of ^d^PMI-δ*(2-6) and* ^d^PMI-δ*(2-9)*; a key hydrogen bond between the backbone of 6-Fluro ^d^Trp at position 3 and the sidechain of Q72 is lost when the residue at position 2 is replaced with a stapled linker as the alpha-methyl interferes with and prevents the formation of this hydrogen bond. Our models are unable to demonstrate whether this is a kinetic effect or a thermodynamic effect; nevertheless, ATSP-7041 is also known to lose affinity when Thr2 is mutated to either Aib or N-methyl Thr [57].

The 1:1 stoichiometric binding was further confirmed by ITC experiments [Figure 5]. Both the linear and stapled ^d^PMI-δ peptides bound to Mdm2 with ΔG = ~ −11 kcal/mol [Supplementary table 1], with the enthalpy of binding (ΔH) ranging from −15.8 kcal/mol for the ^d^PMI-δ*(1-5)* to −8.25 kcal/mol for the ^d^PMI-δ*(5-12).* Both the linear ^d^PMI-δ peptide and ^d^PMI-δ*(1-5)* stapled peptide are less helical in solution, but had a favourable enthalpic contribution to binding. The favourable enthalpy compensates for the entropic penalty paid as the disordered peptide gets ordered during binding to Mdm2. On the other hand, stapled peptide ^d^PMI-δ*(5-12)*, which had increased helicity (34% helicity), makes favourable entropic contributions to the binding, compensating for the loss in favourable enthalpy contributions and therefore binding is retained.

Next the proteolytic stabilities of the stapled and unstapled ^d^PMI-δ peptides were investigated by incubating these molecules in whole cell homogenate and human plasma. As expected, the all-D amino acids composition rendered all peptides (linear and stapled) resistant to proteolytic degradation. Specifically, >90% of each peptide remained detectable in the homogenate during the 22-hour incubation time (Figure S1). Small decreases in peptide concentrations over time were attributed to sample loss due to binding to labware and instrument surfaces rather than through proteolysis. Similarly both the linear and stapled peptides remind stable in the human plasama, with >90% of each peptide remained detectable in the human plasma during the 4-hour incubation time (Figure S1).

### Cellular uptake of stapled D-peptides

To investigate the effect of peptide stapling in the cellular context, linear and stapled ^d^PMI-δ peptides were added to HCT116 cells with a stably transfected p53-responsive β-lactamase reporter gene. After 16 hours of peptide incubation, no p53 activation was observed for the linear ^d^PMI-δ peptide, even at the highest concentration tested (50 µM). In contrast, three of the six stapled ^d^PMI-δ peptides showed dose responsive increases in p53 activity, while the other three were inactive across the range of peptide concentrations tested (Figure 7A).

**Figure 7:**
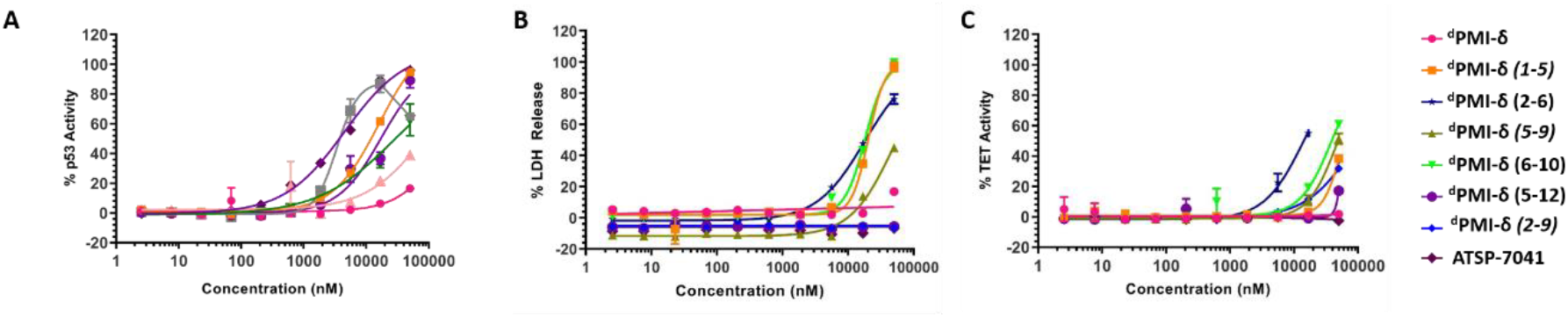
**(A)** stapled ^d^PMI-δ peptides titrated on to HCT116 p53 reporter cells and p53 transcriptional activation assessed in the presence of 2% of serum. (**B)** stapled ^d^PMI-δ peptides titrated on to HCT116 cells and LDH release measured. (**C**) Activity of stapled ^d^PMI-δ peptides measured in a counter screen.

Cellular activity correlated well with the biophysical and biochemical data. Stapled peptides ^d^PMI-δ*(1-5)* and ^d^PMI-δ*(5-12)* bound Mdm2 well (with Kds of 13 and 39 nM respectively from the FP assay) and also demonstrated measurable cellular activation of p53 with EC_50_s of 13.7 and 21.3 µM respectively (Table 1). While these peptides clearly cross the cell membrane in order to activate p53, peptides ^d^PMI-δ*(5-9)*, ^d^PMI-δ*(2-6)* and ^d^PMI-δ*(2-9)* were unable to activate cellular p53, perhaps due their lack of binding affinity (Kd in µM range). Interestingly peptide ^d^PMI-δ*(6-10)* a non-binder of Mdm2 (from the FP assay), demonstrated measurable cellular activation with EC_50_ of 30.3 µM.

### Membrane distribution and counterscreen activity of all-D stapled peptides

Macrocyclic peptides that are cell permeable are often hydrophobic in nature [58], a property that can impart an ability to disrupt the outer membrane and result in cellular leakage [59]. To assess whether the results from our p53 reporter assay were potentially compromised by membrane damage, we carried out a membrane integrity assay (lactate dehydrogenase(LDH) release) under identical conditions to our p53 cellular assay. The linear ^d^PMI-δ peptide which did not show any p53 cellular activity also did not show any LDH leakage at concentrations as high as 50 µM [Figure 7B, Table 1]. The stapled peptides, ^d^PMI-δ*(2-9)* and ^d^PMI-δ*(5-9)* also did not cause LDH leakage, even at concentrations as high as 50 µM. The stapled peptide ^d^PMI-δ(2-9) which was weak binder in biochemical assays and did not result in any cell activity also did not cause LDH release, suggesting that it is cell impermeable. Interestingly the most cell active stapled peptide ^d^PMI-δ*(5-12)*, didn’t cause any LDH leakage, suggesting that the activity observed in the p53 receptor activation assay is through intracellular target engagement. Stapled peptides ^d^PMI-δ*(1-5)*, ^d^PMI-δ*(2-6)* and ^d^PMI-δ*(6-10)* all caused LDH leakage with EC_50_ of 24 µM, 17.7 µM and 18.5 µM respectively. Peptide ^d^PMI-δ*(2-6)*, a non-binder of Mdm2 and without any measurable cell activity, induced LDH leakage with EC_50_ 17.7 µM. Peptides ^d^PMI-δ*(6-10)*, a non-binder of Mdm2 and ^d^PMI-δ*(1-5)* a potent binder of Mdm2 with cell efficacy, also resulted in considerable LDH leakage. In fact, the EC_50_ observed in the p53 reporter activity assays is similar to the EC_50_ observed for the LDH leakage assays, indicating that these two stapled peptides cause membrane disruption and the readout of the p53 reporter assay may not be due to intracellular target engagement but instead due to the nonspecific cytotoxicity resulting from plasma membrane lysis. Thus, we demonstrate that ^d^PMI-δ*(5-12)* enters the cells without membrane disruption, engages the Mdm2-p53 complex, resulting in p53 reporter activity.

To further validate intracellular target engagement, we carried out a counterscreen assay with an identical reporter gene but one whose expression is independent of p53 activation. Most of the stapled ^d^PMI-δ peptides including the linear peptide had EC_50_ values > 50 µM [Figure 7C]. For the linear ^d^PMI-δ peptide and stapled peptides ^d^PMI-δ*(5-9)* and ^d^PMI-δ*(2-9)*, this result was unsurprising as these molecules appear to be cell impermeable. Interestingly, stapled peptides ^d^PMI-δ*(1-5)* and ^d^PMI-δ*(6-10)* each demonstrate significant activity in the reporter assay and LDH leakage assay, and didn’t exhibit activity or were weakly active (^d^PMI-δ*(6-10)* 39 µM) in the counterscreen assay. Stapled peptide ^d^PMI-δ*(2-6)* which had no activity in the reporter assay, demonstrated significant activity in the counterscreen assay and is similar to the EC_50_ observed for the LDH leakage assays. ^d^PMI-δ*(5-12)* displayed negligible cytotoxicity with EC_50_ > 50 µM, suggesting that it is not cytotoxic and acts mechanistically through disruption of the intracellular Mdm2-p53 complex.

### Design, synthesis, binding and cellular activity of double staple and stitched ^d^PMI-δ peptides

Encouraged by the results of the stapled ^d^PMI-δ peptides, with particular interest in ^d^PMI-δ*(5-12)* which showed on-target cellular activity without confounding activities in the LDH release or counterscreen assays, we wondered whether incorporation of an additional staple would confer further improvements in binding and cellular activity. Recent studies have highlighted the limitations of peptides carrying single staples including low cell permeability, low proteolytic stability and low cellular activity and have shown that these can be overcome with the introduction of an additional staple [39, 40, 60]. In such bicyclic arrangements, two pairs of hydrocarbon stapling residues are incorporated into a single peptide sequence. To avoid any cross reactivity during olefin metathesis, sufficient spacing between the two pairs of non-natural amino acid staple precursors are required, often resulting in a longer peptide sequence. Several double-stapled peptides have been shown to successfully inhibit pathways in Rab8a GTP-ase [38], HIV-1 [39], Respiratory Syncytial Virus Entry [40, 60], Ral GTP-ase [61], estrogen receptor-α [62] and BCL9 [63]. All these peptides exhibited increased helicity, increased proteolytic resistance and increased binding as compared to the corresponding single stapled peptides. Some of these double-stapled peptides even demonstrated enhanced cell permeability. Double-stapled peptides can also be designed with a common attachment/anchoring point and peptides with such contiguous hydrocarbon staples are also referred to as “stitched” peptides [64]. Recently Hilinski et al. [64] reported the synthesis of stitched peptides using RCM reactions that exhibited improvements in thermal and chemical stability, proteolytic stability and cell permeability. From the 6 stapled peptides designed we found that two, ^d^PMI-δ*(1-5)* and ^d^PMI-δ*(5-12)*, both exhibited improved binding and cellular properties. We introduced an addition staple between positions 9 and 12 (i+3) in ^d^PMI-δ*(1-5)* resulting in a ^d^PMI-δ*(1-5,9-12)* double stapled peptide [Figure 8]. Combining ^d^PMI-δ*(1-5)* and ^d^PMI-δ*(5-12)* resulted in a stitched peptide, ^d^PMI-δ*(1,5,12)*, with the common attachment point for the two staples localised at residue 5 [Figure 8]. As expected the CD spectra of the stitched ^d^PMI-δ*(1,5,12)* showed increased helicity (52% helicity, Table 2). This agrees with reports on other peptides showing that the stitched and double stapled peptides often display increased helicity compared to the single stapled peptides [38, 39, 40, 60–64]. Curiously, the stitched scrambled peptide had only 28% helicity. In contrast the double stapled peptide ^d^PMI-δ*(1-5,9-12)* did not show enhanced helicity (20 % helicity, Table 2).

**Table 2:** Secondary structure, binding and cellular activity of stapled, double stapled and stitched ^d^PMI-δ peptides determined through various biophysical and biochemical methods.

| Peptide | CD<br>(%helicity) | FP<br>Mdm2<br>Kd (nM) | FP<br>Mdm4<br>Kd (nM) | SPR<br>Kd (nM) | P53 activity<br>EC<sub>50</sub> (μM)<br>(2% FBS,<br>16hrs) | LDH<br>release<br>EC<sub>50</sub> (μM)<br>(2% FBS,<br>16hrs) | Counter<br>screen<br>EC<sub>50</sub> (μM)<br>(2% FBS,<br>16hrs) |
| --- | --- | --- | --- | --- | --- | --- | --- |
| <sup>d</sup>PMI-δ | 20.4 | 36.2 | 43.8 | < 1 | > 50 | > 50 | > 50 |
| <sup>d</sup>PMI-δ(1-5) | 24.7 | 13.2 | 4.5 | < 1 | 13.7 | 24 | > 50 |
| <sup>d</sup>PMI-δ(5-12) | 31.8 | 38.75 | 29.5 | < 1 | 21.3 | > 50 | > 50 |
| <sup>d</sup>PMI-δ(1-5-12) | 52.1 | 4.25 | 12.5 | < 1 | 4 | > 50 | > 50 |
| <sup>d</sup>PMI-δ(1-5-12)<br>Scrambled | 28 | 10000 | 10000 | > 500 | > 50 | > 50 | > 50 |
| <sup>d</sup>PMI-δ(1-5, 9-12) | 20 | 0.7 | 2.1 | < 1 | 34.6 | > 50 | > 50 |
| ATSP-7041 | 49.6 | 1.8 | 4.5 | < 1 | 4.2 | > 50 | > 50 |

**Table 3:**
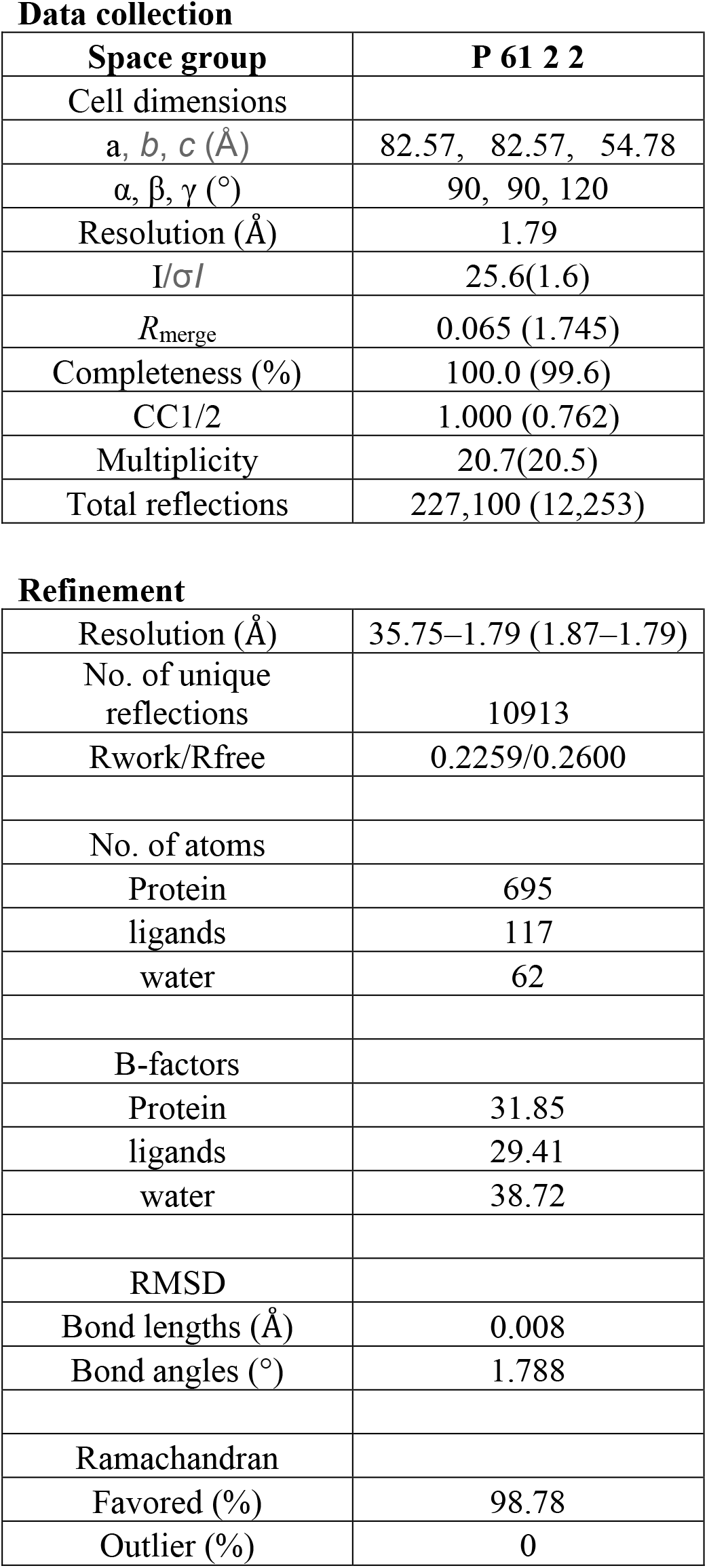
Data collection and refinement Parameters

|  |  |
| --- | --- |
| **Space group** | **P 61 2 2** |
| Cell dimensions |  |
| a, b, c (Å) | 82.57, 82.57, 54.78 |
| $\alpha, \beta, \gamma$ (°) | 90, 90, 120 |
| Resolution (Å) | 1.79 |
| I/σI | 25.6(1.6) |
| $R_{\text{merge}}$ | 0.065 (1.745) |
| Completeness (%) | 100.0 (99.6) |
| CC1/2 | 1.000 (0.762) |
| Multiplicity | 20.7(20.5) |
| Total reflections | 227,100 (12,253) |
| **Refinement** |  |
| Resolution (Å) | 35.75–1.79 (1.87–1.79) |
| No. of unique reflections | 10913 |
| Rwork/Rfree | 0.2259/0.2600 |
| No. of atoms |  |
| Protein | 695 |
| ligands | 117 |
| water | 62 |
| B-factors |  |
| Protein | 31.85 |
| ligands | 29.41 |
| water | 38.72 |
| RMSD |  |
| Bond lengths (Å) | 0.008 |
| Bond angles (°) | 1.788 |
| Ramachandran |  |
| Favored (%) | 98.78 |
| Outlier (%) | 0 |

**Figure 8:**
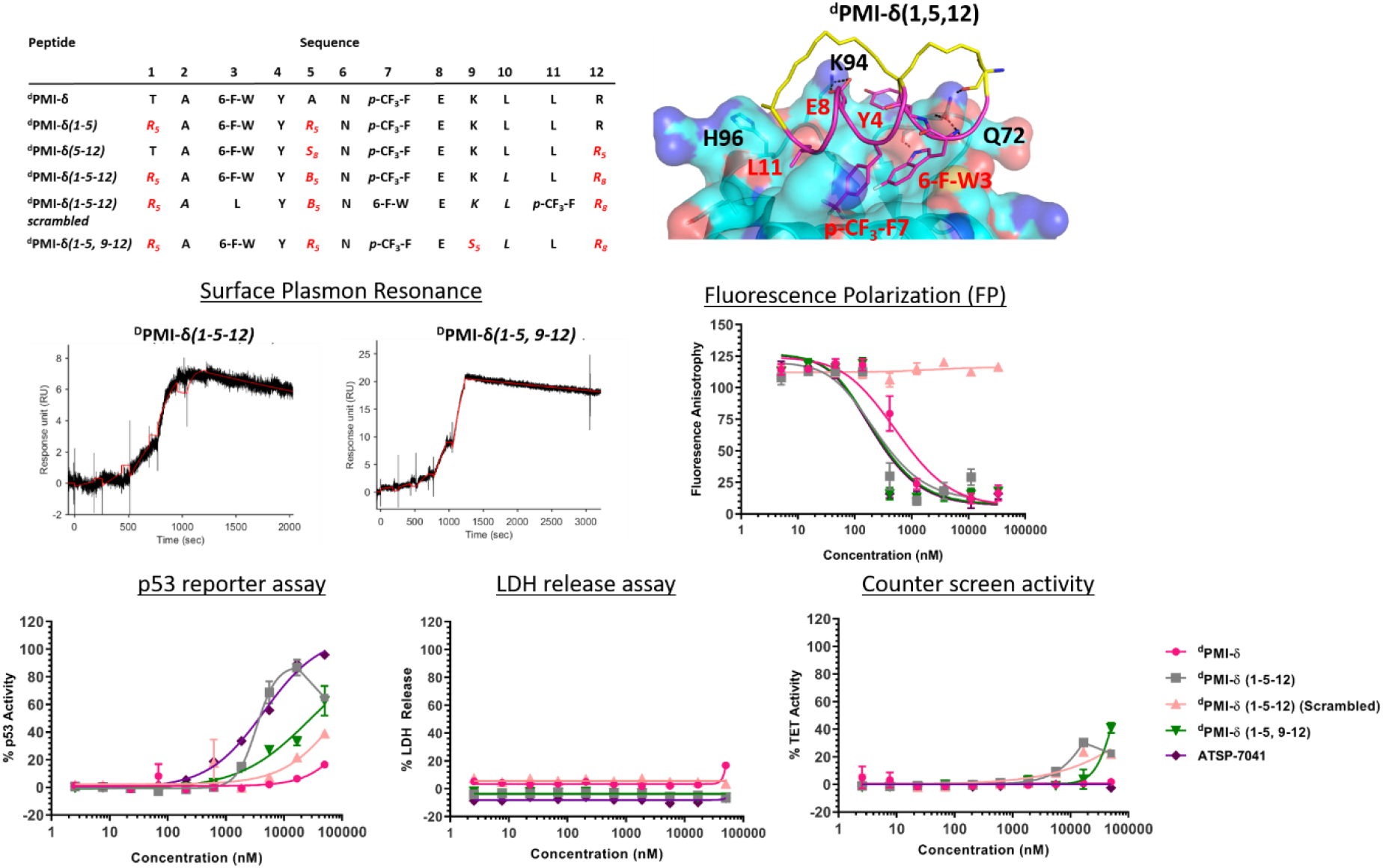
Sequences of stapled and stitched ^d^PMI-δ peptides are shown. MD snapshot of stitched ^d^PMI-δ(1,5,12) -Mdm2 complex. Mdm2 is shown as surface and bound peptide (magenta) is shown as cartoon with interacting residues highlighted in sticks. The hydrocarbon stitch (linker) is highlighted in yellow. Hydrogen bond interactions are shown as dotted lines (black). Binding activity of stapled and stitched ^d^PMI-δ peptides toward Mdm2 protein determined through SPR and Fluorescence polarization assay. Stapled and stitched ^d^PMI-δ peptides were titrated on to HCT116 p53 reporter cells and p53 transcriptional activation assessed in the presence of 2% serum. Stapled and stitched ^d^PMI-δ peptides were titrated on to HCT116 cells and LDH release measured. Activity of stapled and stitched ^d^PMI-δ peptides were measured in a counter screen.

Both the stitched ^d^PMI-δ*(1,5,12)* and double-stapled bicyclic peptides ^d^PMI-δ*(1-5,9-12)* bound Mdm2 with Kd of 4 and 0.7 nM in FP assay, a 10-100 fold increase in affinity as compared to ^d^PMI-δ*(1-5)* and ^d^PMI-δ*(5-12)* [Figure 8, Table 2]. This confirms that the additional staple enhances the target engagement of these peptides. ^d^PMI-δ*(1,5,12)* displayed enhanced cellular activity with EC_50_ of 4 µM [Figure 8, Table 2], at 16 hrs, a five-fold increase compared to the single stapled peptides. Similar increases in cell permeability for a stitched peptide have been reported earlier [64]. The stitched scrambled peptide did not show any binding or cellular activity. However the double-stapled peptide ^d^PMI-δ*(1-5,9-12)* didn’t have significant cellular activity with an EC_50_ of 34.6 µM, at 16 hrs [Figure 8, Table 2]. Although recent studies have reported that the double-stapled peptides appear to follow the same trend as their single-stapled counterparts [38, 39, 40, 60-62], lack of cell activity observed for the double-stapled peptide here, demonstrate that enhanced cellular activity is not uniform. Although the molecular mechanisms behind the increased cell permeability of the stitched D-peptide is unclear, it could be attributed to the increased conformational rigidity and/or increased hydrocarbon content of the peptide. No detectable LDH leakage was observed, even at concentrations as high as 50 µM, and there was negligible counter screen activity, confirming that the designed stitched peptide ^d^PMI-δ*(1,5,12)* enter cells without membrane disruption and result in the activation of p53 by inhibiting the Mdm2-p53 complex [Figure 8, Table 2]. The intracellular target engagement of stitched ^d^PMI-δ*(1,5,12)* peptide was further validated using western-blot analysis. Stabilisation of Mdm2 and activation of p53 was observed for ^d^PMI-δ(1,5,12) and ATSP-7041, whereas the ^d^PMI-δ linear peptide failed to do so [Figure S2].

We also tested the ability of ^d^PMI-δ, ^d^PMI-δ*(1-5,9-12)*, ^d^PMI-δ*(1,5,12)* and scrambled ^d^PMI-δ*(1,5,12)* peptides to inhibit the p53 (1-52)-Mdm2 interaction in a Yeast 2-hybrid (Y2H) assay optimized for detecting small-molecule inhibitors [65]. In this assay, inactivation of p53(1-52)/Mdm2 interaction inhibits growth of yeast cells in a nutrient medium lacking histidine and adenine [65]. Consistent with our observations in mammalian cells, the linear ^d^PMI-δ peptide and the double stapled ^d^PMI-δ*(1-5,9-12)* peptide had no effect on p53-Mdm2 interaction (Figure 9A-B). In contrast, the ^d^PMI-δ*(1,5,12)* peptide completely prevented the growth of yeast cells expressing p53/Mdm2 at 12.5 µM (Figure 9C, left panel). This inhibition was specific as the ^d^PMI-δ*(1,5,12)* peptide had no effect on interaction between two yeast kinetochore proteins Csm1 and Dsn1 (Figure 9C, right panel). Moreover, a scrambled version of ^d^PMI-δ*(1,5,12)* had no effect on p53-Mdm2 interaction (Figure 9D). Our results indicate that the ^d^PMI-δ*(1,5,12)* specifically inhibits p53(1-52)/Mdm2 interaction in yeast cells. Unlike mammalian cells, yeast cells have a thick cell wall. Potency of ^d^PMI-δ*(1,5,12)* in the Y2H assay therefore highlights the ability of stitched peptides to traverse barriers such as the cell wall and plasma membrane.

**Figure 9:**
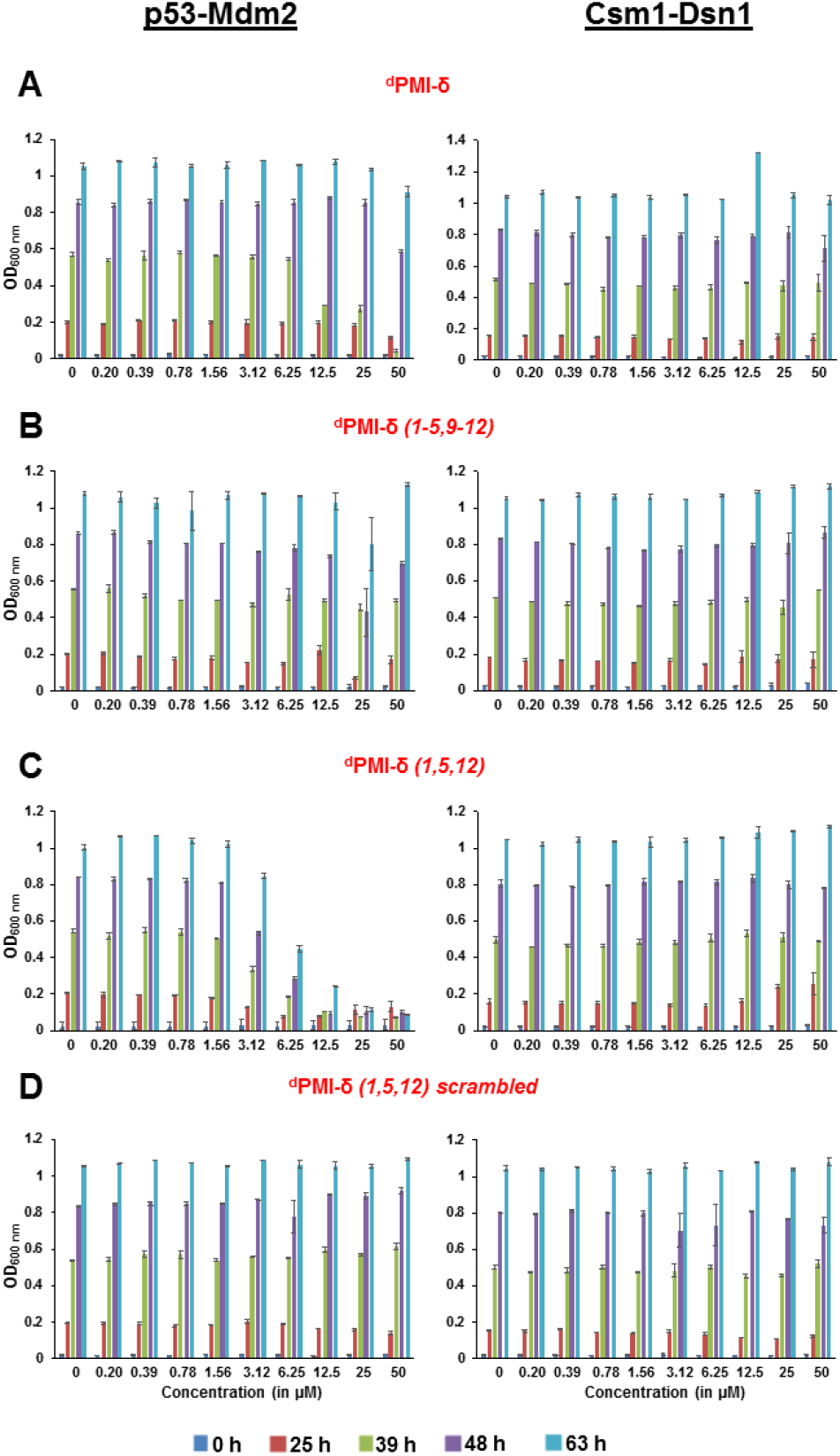
Overnight cultures of ABC9Δ strain containing plasmids encoding either Gal4 AD-p53 (1-52)/Gal4 BD-Mdm2 or Gal4 AD-Csm1/Gal4 BD-Dsn1 were inoculated at OD_600_=0.2 into selective medium containing DMSO or ^d^PMI-δ or ^d^PMI-δ(1-5,9-12), ^d^PMI-δ(1,5,12) and scrambled ^d^PMI-δ(5,12) peptides at the indicated concentrations in duplicate. For each strain, growth as measured by average OD_600_ of duplicate cultures is plotted at different time points following inoculation (0, 25, 39, 48, and 63 h). Ends of the vertical bar indicate the OD_600_ values of the duplicate cultures. Data for the ^d^PMI-δ or ^d^PMI-δ(1-5,9-12), ^d^PMI-δ(1,5,12) and scrambled ^d^PMI-δ(5,12) peptides are shown in panels A, B, C and D respectively. For each peptide, data showing their effect on p53-Mdm2 and Csm1-Dsn1 interactions are presented on the left and right respectively.

### Dual inhibition by ^d^PMI-δ peptides

Mdm4 is homologous to Mdm2 and is also a negative regulator of p53, often found overexpressed in some cancer cells. Studies have shown that dual inhibition (of Mdm2 and Mdm4) appears to be critical for full activation of p53-dependent tumor suppression [34, 35, 37]. Thus, we were interested to know if the all-D peptides had dual-inhibitory properties. As a control, the single stapled peptide ATSP-7041, a validated Mdm2/Mdm4 binder, was observed to bind to Mdm4 with Kd 4.5 nM. The structure of Mdm4 is highly similar to Mdm2, so we generated a model of the ^d^PMI-δ peptide bound to a structure of Mdm4 using the ^d^PMI-δ-Mdm2 structure as template. Models of the stapled/stitched ^d^PMI-δ peptide bound to Mdm4 were generated by incorporating appropriate linkers in the ^d^PMI-δ-Mdm4 structure. We next carried out MD simulations of the (un)stapled/stitched ^d^PMI-δ peptides bound to Mdm4. The stapled peptides remained stable during the MD simulations and remained largely (~95%) helical. The three critical residues 6-F-^d^Trp3, p-CF3-^d^Phe7 and ^d^Leu11 remained buried in the hydrophobic pocket/binding site of Mdm4 [Figure 10]. The hydrocarbon linkers remained largely exposed to solvent without engaging the Mdm4 surface. The binding of ^d^PMI-δ peptides with Mdm4 was further confirmed by FP assay.

**Figure 10:**
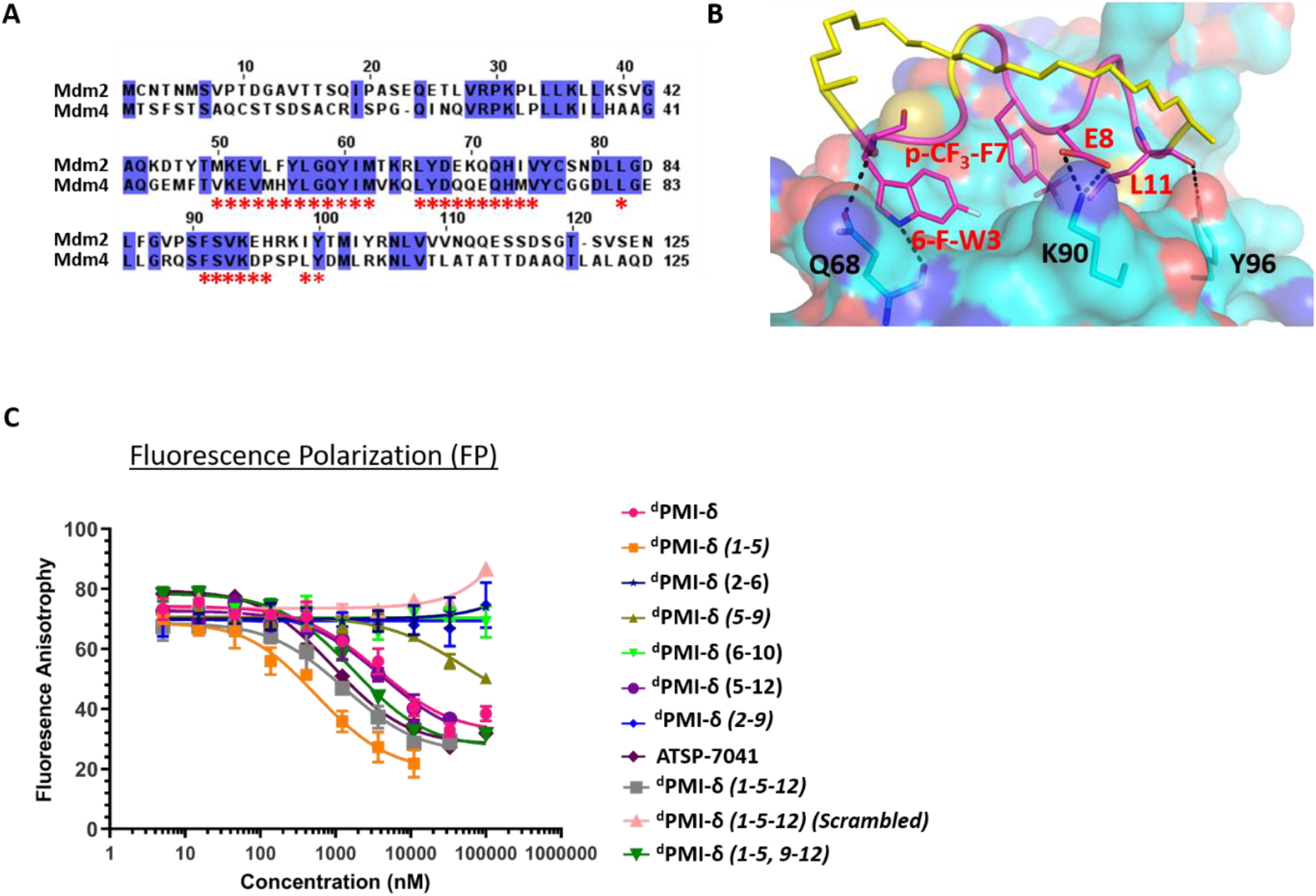
(A) Sequence comparison of N-terminal domains of Mdm2 and Mdm4. Identical residues are highlighted and binding pocket residues (residues that are within 6 Å of bound peptide) are highlighted as *. (B) MD snapshot of stitched ^d^PMI-δ(1,5,12) -Mdm4 complex. Mdm4 is shown as surface and bound peptide is shown as cartoon with interacting residues highlighted as sticks. The hydrocarbon stitch (linker) is highlighted in yellow. Hydrogen bond interactions are shown as dotted lines (black). (C) Fluorescence polarization binding analysis of stapled ^d^PMI-δ peptides and Mdm4 protein.

The Mdm4 binding data mirrored the Mdm2 binding FP assay, with the linear and stapled ^d^PMI-δ(1-5), ^d^PMI-δ(5-12) peptides bind strongly with Kd of 43.8 nM, 4.5 nM and 29.5 nM respectively. The other four stapled peptides ^d^PMI-δ*(2-6)*, ^d^PMI-δ*(5-9)*, ^d^PMI-δ*(2-9)*, ^d^PMI-δ*(6-10)* displayed no binding (Figure 10, Table 1) in the FP assay. Both the stitched ^d^PMI-δ(1,5,12) and double-stapled ^d^PMI-δ(1-5, 9-12) peptides displayed strong affinity towards Mdm4, with Kd of 12.5 and 2.1 nM respectively (Figure 10, Table 2). In conclusion, the stitched ^d^PMI-δ(1,5,12) and double-stapled ^d^PMI-δ(1-5, 9-12) peptides are high affinity dual inhibitors of Mdm2 and Mdm4.

### Crystal structure of Mdm2 - ^d^PMI-δ(1-5, 9-12) complex

We tried to obtain crystal structures of the stitched peptide ^d^PMI-δ(1,5,12) in complex with Mdm2 (6-125) and of the double stapled peptide ^d^PMI-δ(1-5, 9-12) in complex with Mdm2 (6-125). We could only obtain crystals for the latter and were able to resolve its structure (Figures 11 A, B). The elucidated structure possessed only a single copy of the Mdm2 - ^d^PMI-δ(1-5, 9-12) complex in the asymmetric unit. The p53 peptide binding groove on Mdm2 was occupied by a single molecule of ^d^PMI-δ(1-5, 9-12), which interacted with Mdm2 by projecting the following three residues 6-F-^d^Trp3, p-CF3-^d^Phe7 and ^d^Leu11 deep into the hydrophobic peptide binding site on Mdm2. The Mdm2 - ^d^PMI-δ(1-5, 9-12) complex was overlaid with the Mdm2 - ^d^PMI-δ structure (PDB: ID 3TPX), which demonstrated that the double stapling maintained the overall fold of the bound peptide and its critical interactions with Mdm2 [Figure 11 C]. The computationally modelled structure of the Mdm2 - ^d^PMI-δ(1-5, 9-12) complex is very similar to the co-crystal structure of the Mdm2 - ^d^PMI-δ(1-5, 9-12) complex; the peptide rmsd is < 1Å between the two, and the overall fold along with the protein-peptide interactions is in good agreement [Figure 11 D]. Further, as predicted, neither hydrocarbon staple linker engage in any contacts with the surface of Mdm2 and both are exposed into the solvent.

**Figure 11:**
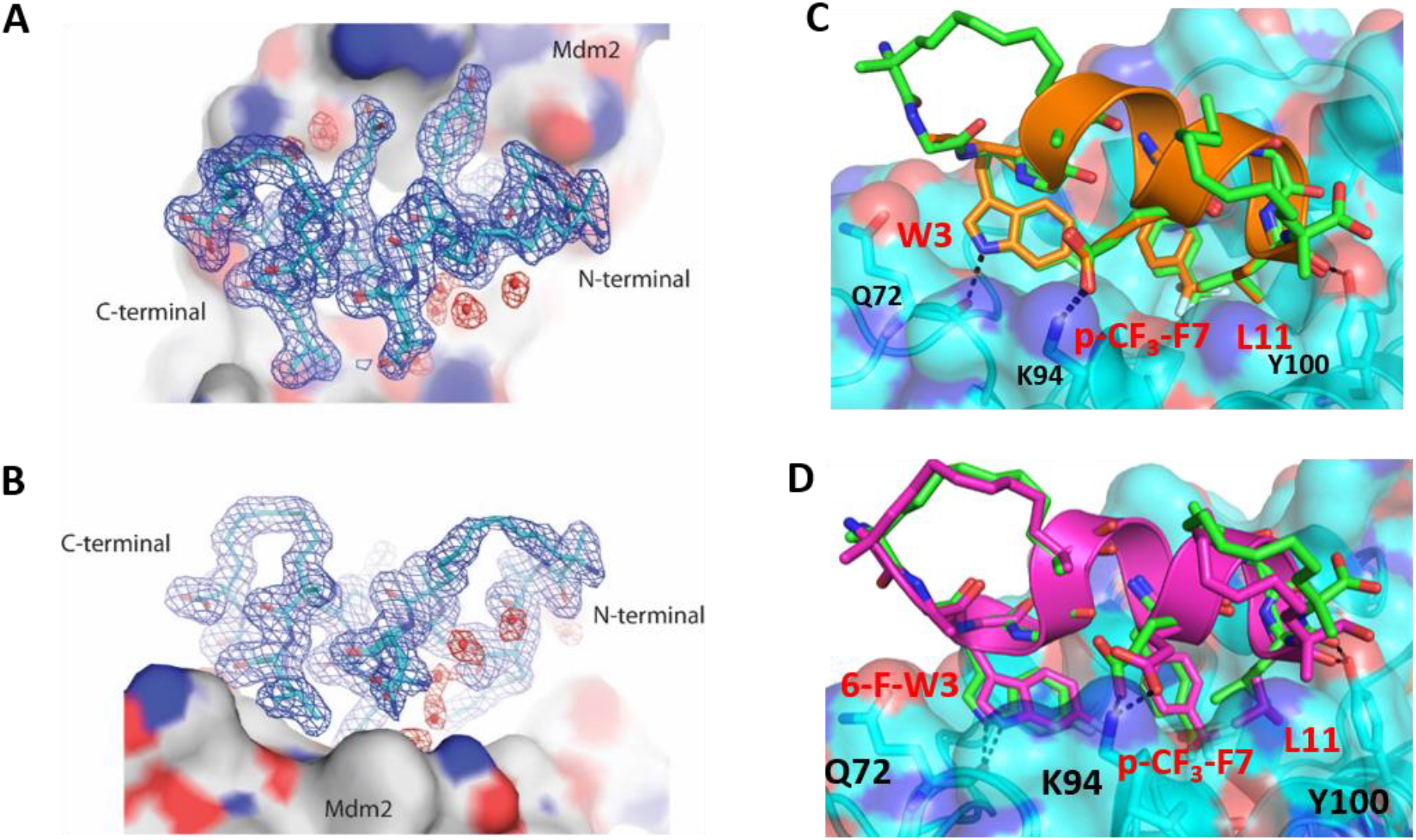
(A) Top down view of ^d^PMI-δ (1-5, 9-12) bound to Mdm2. (**B)** 90° rotation of the ^d^PMI-δ (1-5, 9-12)-Mdm2 complex orientation shown in panel A), which clearly depicts the presence of both hydro-carbon staples in ^d^PMI-δ (1-5, 9-12). The 2F_o_-F_c_ map electron density map (shown with the blue mesh) for ^d^PMI-δ (1-5, 9-12) was contoured at 1.0σ. Interfacial structured waters involved in Mdm2-^d^PMI-δ (1-5, 9-12) complex formation are shown using the red mesh. **(C)** Comparison of crystal structure of ^d^PMI-δ (1-5, 9-12) - Mdm2 complex with the ^d^PMI-δ-Mdm2 structure (PDB: ID 3TPX). The Mdm2 is shown as surface (cyan) with the bound ^d^PMI-δ (1-5, 9-12) (green) and ^d^PMI-δ (orange) peptides are shown as cartoon. The three critical residues (W3, p-CF3-F7 and L11 in ^d^PMI, 6-F-W3, p-CF3-F7 and L11 in ^d^PMI-δ (1-5, 9-12)) and residues involved in protein – peptide interactions are shown as sticks and the interactions are highlighted (black dashed lines). **(D)** Comparison of computationally modelled structure of ^d^PMI-δ (1-5, 9-12) - Mdm2 complex with the crystal structure of ^d^PMI-δ (1-5, 9-12) - Mdm2 complex. The Mdm2 is shown as surface (cyan) with the bound modelled ^d^PMI-δ (1-5, 9-12) (magenta) and experimental ^d^PMI-δ (1-5, 9-12) (green) peptides are shown as cartoon. The three critical residues (6-F-W3, p-CF3-F7 and L11) and residues involved in protein – peptide interactions are shown as sticks and the interactions are highlighted (black dashed lines).

## Discussion

Peptide based inhibitors are emerging as next generation therapeutic modalities because of their high target specificity, high biocompatibility and low toxicity. However, liabilities such as conformational instability, proteolytic sensitivity and lack of cell permeability hinder their potential. Peptide stapling to constrain peptides in their active/bound conformations results in several benefits such as improved stability, target binding and cell permeability. Although stapling L-amino acid peptides can confer resistance to protease-mediated degradation, the effect is often not complete, especially for residues located outside of the staple. On the other hand, D-amino acid peptides show complete resistance to proteolysis, increased stability and bioavailability, hence appear to be suitable for oral administration. Unfortunately, just like most linear peptides, all-D peptides generally lack membrane permeability and cellular activity.

We speculated that a combination of the two strategies (*i.e.*, all D-amino acids and stapling) might provide a robust molecule satisfying all the required criteria for intracellular target engagement. Accordingly, we embarked on introducing a hydrocarbon staple into an all-D peptide inhibitor of the p53-Mdm2/Mdm4 interaction that had been discovered using mirror – image phage display [41]. Guided by the available crystal structure of ^d^PMI-δ bound to the N-terminal domain of Mdm2, we designed six stapled ^d^PMI-δ peptides using a combination of modelling and molecular simulations. All 6 stapled peptides displayed helicity ranging from 24% to 40% which compared with ~21% for the linear counterpart. Two of the peptides demonstrated increased affinity for Mdm2 in biophysical and biochemical experiments. These peptides also showed enhanced cell uptake without detectable membrane disruption and disrupted the Mdm2-p53 complex, leading to activation of p53. No correlation was apparent between helicity and binding or with cell activity as has also been reported for L-amino acid peptides.

We next decided to introduce a second staple generating a double stapled peptide and a stitched peptide. The stitched peptide displayed the highest helicity (52%) while the double stapled peptide remained unchanged at 20%, similar to that adopted by the linear peptide. Nevertheless, both these peptides displayed increased binding to Mdm2, suggesting that the binding mechanism of these peptides are different from each other. However only the stitched peptide displayed increased cellular activity, probably resulting from increased cell permeability; the double stapled peptide appears unable to cross the cell membrane. Although stitching resulted in increased helicity, increased affinity and more importantly enables cell permeability, it is not clear how this latter is achieved. Increased hydrophobicity resulting from the hydrocarbon linker of the stitched peptide could be a major driving force, however the lack of permeability for the double stapled peptide which has a longer hydrophobic linker casts doubt on the hydrophobicity – permeability link. Therefore, understanding cell permeability warrants further studies to systematically investigate the factors enabling cell permeability of these peptides.

While stapling appeared to impart cell permeability, some of the stapled peptides caused membrane disruption. Curiously, while all the stapled and stitched peptides displayed reporter activity at 4 hrs, only 4 peptides (^d^PMI-δ(1-5), ^d^PMI-δ(5-12), ^d^PMI-δ(6-10), ^d^PMI-δ(1-5-12)) continued to show activity at 16 hours. However, ^d^PMI-δ(6-10) is known to not bind Mdm2 and is also known to cause membrane disruption (from LDH release assays) and hence it likely activate p53 due to membrane disruption related activation of stress pathways and not through direct engagement with Mdm2. At the same time, ^d^PMI-δ(1-5), which is a potent binder of Mdm2, also causes LDH leakage and hence it is unclear what results in p53 activation: target engagement or membrane disruption, likely some combination of the two.

A counterscreen assay, with an identical read out to our primary cellular screen but that is independent of p53 activation, was carried out to find peptides with off-target effects. We found that the peptides ^d^PMI-δ(2-6) and ^d^PMI-δ(6-10) both non-binders of Mdm2 and membrane disrupters, activated p53 even at 16 hours, showed counter screen activity; this could result from membrane disruption or off-target activity. In contrast the ^d^PMI-δ(5-12) and the stitched peptide showed no membrane disruption and off target activity, activating p53 through intracellular target engagement. The on-target engagement of stapled and stitched peptide was further validated in western blot assays. Thus, it is important to use a combination of LDH leakage, counter screen assays and target engagement/reporter activation to rule out false positives (that result from off-target engagement and membrane disruption). The yeast assay confirmed the on target activity of the ^d^PMI-δ(1-5-12) peptide adding support to the favourable effects of this approach to peptide stapling on cell permeability.

While we were unable to obtain crystals of the stitched peptide in complex with Mdm2, we successfully resolved the structure of the double stapled peptide in complex with Mdm2. The latter showed that the critical interactions of the peptide with Mdm2 were retained and the hydrocarbon staple linker points into solvent, without engaging in any contacts with Mdm2. Given that our computationally predicted structure matched the crystal structure to within 1 Å, we are confident that our model of the stitched peptide complexed to Mdm2 would also closely mirror the actual complex; the key interactions with Mdm2 are maintained and the staple points into solvent without engaging in any contacts with Mdm2 (in any case the 1-5 staple is common to both structures).

Stapling also enabled the peptide to bind to Mdm4 with high affinity, The Mdm4 binding data mirrored the Mdm2 data, resulting in a cell permeable dual inhibitor of Mdm2/Mdm4. It is possible that in the activation assays, binding to Mdm4 likely contributes to p53 activation. Several studies have shown that dual inhibition of Mdm2 and Mdm4 appears to be critical for full activation of p53-dependent tumour suppression.

In conclusion, by stapling all D-amino acid peptides, we have successfully demonstrated that stapling can enhance both binding and cellular properties of linear peptides, as has been reported for the L-amino acid peptides. The use of all-D peptides leveraging intrinsic stability and macrocyclization as described here imparts enhanced target binding and cellular activity to advance a novel stapled peptide modality having significant therapeutic potential for p53-dependent cancers.

## Supporting Information

- Full Computational and experimental details
- Whole cell homogenate and human plasma stability results
- Western-blot analysis

## Acknowledgement

We would like to acknowledge Yan Lin and Hochman Jerome for critical reading of the manuscript. The authors thank National Super Computing Centre (NSCC) for computing facilities. This work was supported by a collaborative grant from MSD to A*STAR (p53lab, ICES, and BII) and an Industry Alignment Pre-Positioning Grant (Peptide Engineering Program) to A*STAR (p53lab, ICES, and BII) (grant IDs H17/01/a0/010, IAF111213C).

