## Supplementary Information for "Macrocyclization of an all-D linear peptide improves target affinity and imparts cellular activity: A novel stapled α-helical peptide modality"

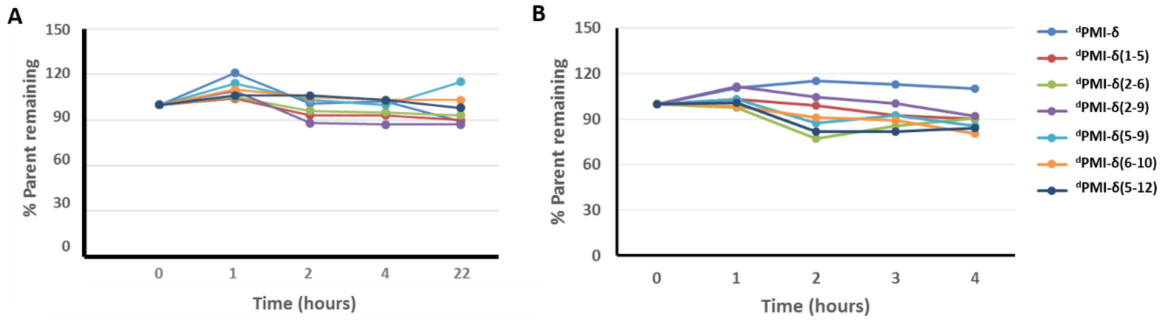


**Figure S1:** (A) Stability of stapled and linear $^{d}$PMI-δ peptides in whole cell homogenate. Parent sequences. (B) Plasma stability of the stapled $^{d}$PMI-δ peptides quantified over 4 hrs.

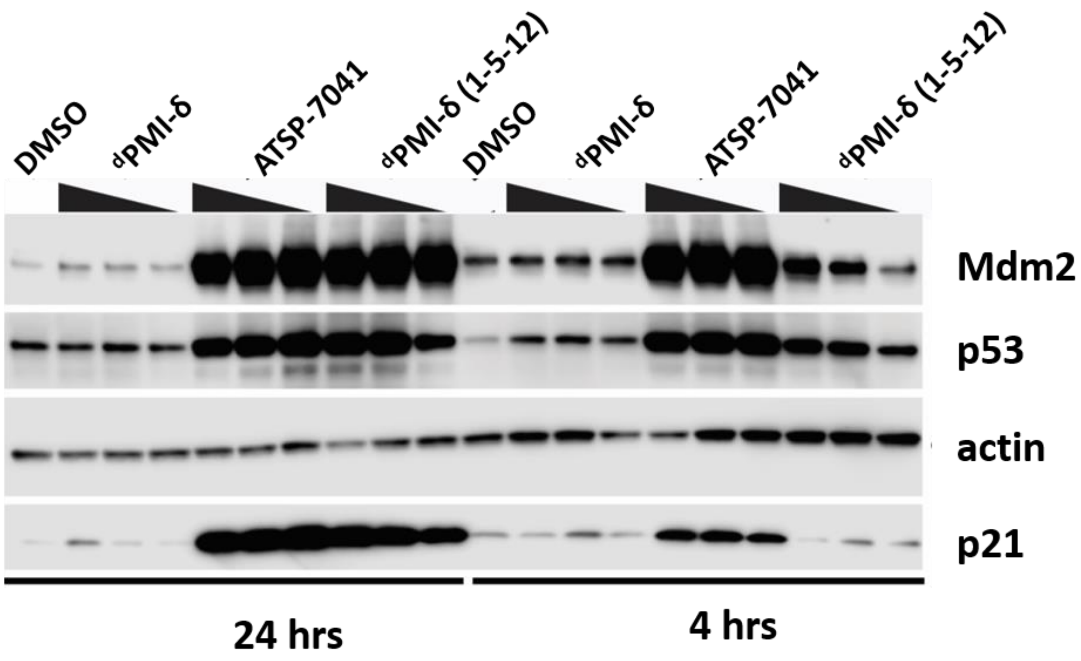


**Figure S2:** Western blot analysis of HCT-116 cells treated with either vehicle control (1% DMSO) or with 6.123 µM, 12.5 µM and 25 µM of the stated compound for either 4 or 24 hours. Compounds treatments contained a residual DMSO concertation of 1% v/v DMSO**.**

**Supplementary table1:** Binding of stapled & stitched $^{d}$PMI-δ peptides toward MDM2 protein determined through ITC experiments.

| Peptide | ITC (kcal/mol) | | |
|---|---|---|---|
| | ΔH | -TΔS | ΔG |
| $^{d}$PMI-δ | -13.80 | 3.04 | -10.8 |
| $^{d}$PMI-δ*(1-5)* | -15.8 | 4.2 | -11.6 |
| $^{d}$PMI-δ*(2-6)* | n.d | n.d | n.d |
| $^{d}$PMI-δ*(2-9)* | n.d | n.d | n.d |
| $^{d}$PMI-δ*(5-9)* | n.d | n.d | n.d |
| $^{d}$PMI-δ*E(5-12)* | -8.25 | -1.83 | -10.1 |
| $^{d}$PMI-δ*(6-10)* | -9.35 | -1.86 | -11.2 |

### Materials and Methods

#### Computational

The available crystal structure of the linear dPMI ($^{d}$PMI-δ) peptide co-crystalized with Mdm2 [pdb 3TPX] [1] was used to model the stapled (single and double) and stitched peptides. All models were subjected to molecular dynamics (MD) simulations for further refinement. MD simulations were carried out on free peptide and also on the peptide – Mdm2 complexes. The Xleap module of the Amber18 program [2] was used to prepare the system for the MD simulations. Hydrogen atoms were added and the C- terminus of the peptide was capped with the NHE moiety. The parameters for the staple linkers were taken from our previous study [3]. All simulation systems were neutralized with an appropriate number of counter ions. The neutralized system was solvated in an octahedral box with TIP3P [4] water molecules, leaving at least 10 Å between the solute atoms and the borders of the box. MD simulations were carried out with the pmemd module of the Amber18 package in combination with the ff14SB force field [5]. All MD simulations were carried out in explicit solvent at 300 K. During all the simulations the long-range electrostatic interactions were treated with the particle mesh Ewald [6] method using a real space cut off distance of 9 Å. The settle [7] algorithm was used to constrain bond vibrations involving hydrogen atoms, which allowed a time step of 2 fs during the simulations. Solvent molecules and counter ions were initially relaxed using energy minimization with restraints on the protein and peptide atoms. This was followed by unrestrained energy minimization to remove any steric clashes. Subsequently the system was gradually heated from 0 to 300 K using MD simulations with positional restraints (force constant: 50 kcal $mol^{-1}$ $Å^{-2}$) on protein and peptides over a period of 0.25 ns allowing water molecules and ions to move freely followed by gradual removal of the positional restraints and a 2 ns unrestrained equilibration at 300 K. The resulting systems were used as starting structures for the respective production phase of the MD simulations. For each case, three independent (using different initial random velocities) MD simulations were carried out starting from the well equilibrated structures. Each MD simulation was carried out for 250 ns and conformations were recorded every 4 ps. To enhance the conformational sampling, each of these peptides were subjected to Biasing Potential Replica Exchange MD (BP-REMD) simulations. The BP-REMD technique is a type of Hamiltonian -REMD methods which includes a biasing potential that promotes dihedral transitions along the replicas [8, 9]. For each system, BP-REMD was carried out with eight replicas including a reference replica without any bias. BP-REMD was carried out for 50 ns with exchange between the neighbouring replicas attempted for every 2 ps and accepted or rejected according to the metropolis criteria. Conformations sampled at the reference replica (no bias) was used for further analysis. Simulation trajectories were visualized using VMD [10] and figures were generated using Pymol [11].

**Binding Energy calculations and energy decomposition analysis**

Molecular Mechanics Poisson Boltzmann Surface Area (MMPBSA) methods were used for the calculation of binding free energies between the peptides and their partner proteins [12, 13] 250 conformations extracted from the last 50 ns of the simulations were used for the binding energy calculations. Entropy calculations are computationally intensive and do not converge easily and hence are ignored. The effective binding energies were decomposed into contributions of individual residues using the MMGBSA energy decomposition scheme. The MMGBSA calculations were carried out in the same way as in the MMPBSA calculations. The polar contribution to the solvation free energy was determined by applying the generalized born (GB) method (igb =2) [2], using mbondi2 radii. The non-polar contributions were estimated using the ICOSA method [2] by a solvent accessible surface area (SASA) dependent term using a surface tension proportionality constant of 0.0072 kcal/mol $Å^2$. The contribution of peptide residues was additionally explored by carrying out in-silico alanine scanning in which each peptide residue is mutated to D-alanine in each conformation of the MD simulation and the change with respect to the binding energy of the wild type peptide is calculated using MMPBSA and averaged over all the conformations.

**Peptide Synthesis**

Peptides were synthesized using Rink Amide MBHA resin and Fmoc-protected amino acids, coupled sequentially with DIC/HOBt activating agents. Double coupling reactions were performed on the first amino acid and also at the stapling positions. At these latter positions, the activating reagents were switched to DIEA/HATU for better coupling efficiencies. Ring closing metathesis reactions were performed by first washing the resin three times with DCM, followed by the addition of the first-generation Grubbs Catalyst (20 mol % in DCM and allowed to react for 2 h; all steps with Grubbs Catalyst were performed in the dark). The RCM (ring closing metathesis) reaction was repeated to ensure a complete reaction. After the RCM was complete, a test cleavage was performed to ensure adequate yield. Peptides were cleaved and then purified as a mixture of cis-trans isomers by RP-HPLC.

**Mdm2 Protein Production**

For use in the peptide binding assay, a human Mdm2 1–125 sequence was cloned into a pNIC-GST vector. The TEV (tobacco etch virus) cleavage site was changed from ENLYFQS to ENLYFQG to give a fusion protein with the following sequence:

MSDKIIHSPILGYWKIKGLVQPTRLLLEYLEEKYEEHLYERDEGDKWRNKKFELGLE
FPNLPYYIDGDVKLTQSMAIIRYIADKHNMLGGCPKERAEISMLEGAVLDIRYGVSRI
AYSKDFETLKVDFLSKLPEMLKMFEDRLCHKTYLNGDHVTHPDFMLYDALDVVLY
MDPMCLDAFPKLVCFKKRIEAIPQIDKYLKSSKYIAWPLQGWQATFGGGDHPPKLEV
LFQGHMHHHHHHSSGVDLGTENLYFQGMCNTNMSVPTDGAVTTSQIPASEQETLVR
PKPLLLKLLKSVGAQKDTYTMKEVLFYLGQYIMTKRLYDEKQQHIVYCSNDLLGDL
FGVPSFSVKEHRKIYTMIYRNLVVVNQQESSDSGTSVSEN-.

The corresponding plasmid was transformed into BL21 (DE3) Rosetta T1R Escherichia coli cells and grown under kanamycin selection. Bottles of 750 mL Terrific Broth, supplemented with appropriate antibiotics and 100 μL of antifoam 204 (Sigma-Aldrich, St. Louis, MO, USA, were inoculated with 20 mL seed cultures grown overnight. The cultures were incubated at 37 °C in the LEX system (Harbinger Biotech, Toronto, Canada) with aeration and agitation through the bubbling of filtered air through the cultures. The LEX system temperature was reduced to 18 °C when culture OD600 reached 2, and the cultures were induced after 60 min with 0.5 mM IPTG. Protein expression was allowed to continue overnight. Cells were harvested by centrifugation at 4000× g, at 15 °C for 10 min. The supernatants were discarded and the cell pellets were resuspended in a lysis buffer (1.5 mL per gram of cell pellet). The cell suspensions were stored at −80 °C before purification work.

The re-suspended cell pellet suspensions were thawed and sonicated (Sonics Vibra-Cell, Newtown, CO, USA) at 70% amplitude, 3 s on/off for 3 min, on ice. The lysate was clarified by centrifugation at 47,000× g, 4 °C for 25 min. The supernatants were filtered through 1.2 μm syringe filters and loaded onto the AKTA Xpress system (GE Healthcare, Fairfield, CO, USA). The purification regime is briefly described as follows. The lysates were loaded onto a 1 mL Ni-NTA Superflow column (Qiagen, Valencia, CA, USA) that had been equilibrated with 10 column volumes of wash 1 buffer. Overall buffer conditions were as follows: Immobilized metal affinity chromatography (IMAC) wash 1 buffer—20 mM HEPES ((4-(2-hydroxyethyl)-1-piperazineethanesulfonic acid), 500 mM NaCl, 10 mM Imidazole, 10% (v/v) glycerol, 0.5 mM TCEP (Tris(2-carboxyethyl)phosphine), pH 7.5; IMAC wash 2 buffer—20 mM HEPES,

500 mM NaCl, 25 mM Imidazole, 10% (v/v) glycerol, 0.5 mM TCEP, pH 7.5; IMAC Elution buffer—20 mM HEPES, 500 mM NaCl, 500 mM Imidazole, 10% (v/v) glycerol, 0.5 mM TCEP, pH 7.5. The sample was loaded until air was detected by the air sensor, 0.8 mL/min. The column was then washed with wash 1 buffer for 20 column volumes, followed by 20 column volumes of wash 2 buffer. The protein was eluted with five column volumes of elution buffer. The eluted proteins were collected and stored in sample loops on the system and then injected into gel filtration (GF) columns. Elution peaks were collected in 2 mL fractions and analyzed on SDS-PAGE gels. The entire purification was performed at 4 °C. Relevant peaks were pooled, TCEP was added to a total concentration of 2 mM. The protein sample was concentrated in Vivaspin 20 filter concentrators (VivaScience, Littleton, MA, USA) at 15 °C to approximately 15 mg/mL. (<18 kDa—5 K MWCO, 19–49 kDa—10 K MWCO, >50 kDa—30 K MWCO). The final protein concentration was assessed by measuring absorbance at 280 nm on Nanodrop ND-1000 (Thermo Fisher, Waltham, MA, USA). The final protein purity was assessed on SDS-PAGE gel. The final protein batch was then aliquoted into smaller fractions, frozen in liquid nitrogen and stored at −80 °C.

For x-ray crystallography, Mdm2 (6–125) was cloned as a GST-fusion protein using the pGEX-6P-1 GST expression vector (GE Healthcare). The GST-fused Mdm2 (6–125) construct was then transformed into Escherichia coli BL21(DE3) pLysS (Thermo Fisher, Waltham, MA, USA) competent cells. Cells were grown in Luria-Bertani (LB) medium at 37 °C and induced at OD600 nm of 0.6 with 0.5 mM Isopropyl β- D -1-thiogalactopyranoside (IPTG) at 16 °C. After overnight induction, the cells were harvested by centrifugation, resuspended in binding buffer (50 mM Tris-HCl pH 8.0, 150 mM NaCl), and lysed by sonication. After centrifugation for 60 min at 19,000× g at 4 °C, the cell lysate was then applied to a 5 mL GSTrap FF column (GE Healthcare) pre-equilibrated in wash buffer (50 mM Tris-HCl pH 8.0, 150 mM NaCl, 1 mM DTT). The GST-fused Mdm2 (6–125) was then cleaved on-column by PreScission protease (GE Healthcare) overnight at 4 °C and eluted off the column with wash buffer. The protein sample was then dialyzed into a buffer A solution (20 mM Bis-Tris, pH 6.5, 1 mM DTT) using HiPrep 26/10 Desalting column, and loaded onto a cation-exchange Resource S 1 mL column (GE Healthcare), pre-equilibrated in buffer A. The column was then washed in six column volumes of buffer A and the bound protein was eluted with a linear gradient in buffer comprising 1 M NaCl, 20 mM Bis-Tris pH 6.5, and 1 mM DTT over 30 column volumes. Protein purity as assessed by SDS-PAGE was ~95%, and the proteins were concentrated using

Amicon-Ultra (3 kDa MWCO) concentrator (Millipore, Burlington, MA, USA). Protein concentration was determined using 280 nm absorbance measurements.

#### Mdm4 protein production

MDM4 protein was cloned into pNIC-GST vector and expressed in LEX system (Harbinger Biotech) at Protein Production Platform (PPP) at NTU School of Biological Sciences. Using glycerol stocks, inoculation cultures were started in 20mL Terrific Broth with 8g/L glycerol supplemented with Kanamycin. The cultures were incubated at 37 °C, 200rpm overnight. The following morning, bottles of 750mL Terrific Broth with 8g/L glycerol supplemented with Kanamycin and 100uL of antifoam 204 (Sigma-Aldrich) were inoculated with the cultures. The cultures were incubated at 37 °C in the LEX system with aeration and agitation through the bubbling of filtered air through the cultures. When the OD600 reached ~2, the temperature was reduced to 18 °C and the cultures were induced after 30 to 60 min with 0.5mM IPTG. Protein expression was allowed to continue overnight. The following morning, cells were harvested by centrifugation at 4200 rpm at 15 °C for 10min. The supernatants were discarded and the cells were re-suspended in lysis buffer (100 mM HEPES, 500 mM NaCl, 10 mM Imidazole, 10 % glycerol, 0.5 mM TCEP, pH 8.0 with Benzonase (4uL per 750mL cultivation) and 250U/uL Merck Protease Inhibitor Cocktail Set III, EDTA free (1000x dilution in lysis buffer) from Calbiochem) at 200 rpm, 4 °C for approximately 30min and stored at -80 °C. The re-suspended cell pellet suspensions were thawed and sonicated (Sonics Vibra-cell) at 70 % amplitude, 3s on/off for 3min, on ice. The lysate was clarified by centrifugation at 47000g, 4 °C for 25min. The supernatants were filtered through 1.2 µm syringe filters and loaded onto AKTA Xpress system (GE Healthcare) with a 1mL Ni-NTA Superflow (Qiagen) IMAC column. The column was washed with 20 column volume (CV) of wash buffer 1 (20 mM HEPES, 500 mM NaCl, 10 mM Imidazole, 10 % (v/v) glycerol, 0.5 mM TCEP, pH 7.5) and 20 CV of wash buffer 2 (20 mM HEPES, 500 mM NaCl, 25 mM Imidazole, 10 % (v/v) glycerol, 0.5 mM TCEP, pH 7.5) or until a stable baseline for 3 min and delta base 5mAU (0.8mL/min) was obtained respectively. Mdm4 protein was eluted with elution buffer (20 mM HEPES, 500 mM NaCl, 500 mM Imidazole, 10 % (v/v) glycerol, 0.5 mM TCEP, pH 7.5) and eluted peaks (start collection: >50mAU, slope >200mAU/min, stop collection: <50mAU, stable plateau of 0.5min, delta plateau 5mAU) were collected and stored in sample loops on the system and then injected into equilibrated Gel Filtration (GF) column (HiLoad 16/60 Superdex 200 prep grade

(GE Healthcare)) and eluted with 20 mM HEPES, 300 mM NaCl, 10% (v/v) glycerol, 0.5 mM TCEP, pH 7.5 at a flowrate of 1.2mL/min. Elution peaks (start collection: >20mAU, slope >10mAU/min, stop collection: < 20mAU, slope >10mAU/min, minimum peak width 0.5min) were collected in 2 mL fractions. The entire purification was performed at 4 °C. Relevant peaks were pooled and TCEP was added to a final concentration of 2 mM. The protein sample was concentrated in Vivaspin 20 filter concentrators (VivaScience) at 15 °C to approximately 15mg/mL. The final protein concentration was assessed by measuring absorbance at 280nm on Nanodrop ND-1000 (Nano-Drop Technologies). The final protein purity was assessed by SDS-PAGE and purified MDM4 protein was frozen in liquid nitrogen and stored at -80 °C.

**Competitive Fluorescence Anisotropy Assays (Mdm2 and Mdm4)**

Purified Mdm2 (1-125) protein was titrated against 50 nM carboxyfluorescein (FAM)-labeled 12/1 peptide13 (FAM-RFMDYWEGL-NH2). Dissociation constants for titrations of Mdm2 and Mdm4 against FAM-labeled 12/1 peptide were determined by fitting the experimental data to a 1:1 binding model equation shown below:

Equation 1:

$$r = r_o + (r_b - r_o) \times \frac{(K_d + [L]_t + [P]_t) - \sqrt{(K_d + [L]_t + [P]_t)^2 - 4[L]_t[P]_t}}{2[L]_t}$$

[P] is the protein concentration (Mdm2), [L] is the labeled peptide concentration, r is the anisotropy measured, $r_0$ is the anisotropy of the free peptide, $r_b$ is the anisotropy of the Mdm2–FAM-labeled peptide complex, $K_d$ is the dissociation constant, $[L]_t$ is the total FAM labeled peptide concentration, and $[P]_t$ is the total Mdm2 concentration. The apparent $K_d$ values for FAM-labeled 12/1 peptide against Mdm2 and Mdm4 were determined to be 13.0 nM and 4.0 nM, respectively. These values were then used to determine apparent $K_d$ values of the respective competing ligands in subsequent competition assays in fluorescence anisotropy experiments.

Mdm2 and Mdm4 competition experiments were performed with their respective concentrations held constant at 250 nM and 75 nM, in the presence of 50 nM of FAM-labeled 12/1. The competing molecules were then titrated against the complex of the FAM-labeled peptide and protein. Apparent Kd values were determined by fitting the experimental data to the equations shown below:

$$r = r_o + (r_b + r_o) \times \frac{2\sqrt{(d^2 - 3e)}\cos(\theta/3) - 9}{3K_{d1} + 2\sqrt{(d^2 - 3e)}\cos(\theta/3) - d}$$

$$d = K_{d1} + K_{d2} + [L]_{st} + [L]_t - [P]_t$$

$$e = ([L]_t - [P]_t)K_{d1} + ([L]_{st} - [P]_t)K_{d2} + K_{d1}K_{d2}$$

$$f = -K_{d1}K_{d2}[P]_t$$

$$\theta = ar\cos\left[\frac{-2d^3 + 9de - 27f}{2\sqrt{(d^2 - 3e)^3}}\right]$$

$[L]_{st}$ and $[L]_t$ denote labeled ligand and total unlabeled ligand input concentrations, respectively. $K_{d2}$ is the dissociation constant of the interaction between the unlabeled ligand and the protein. In all competition experiments, it is assumed that $[P]t > [L]_{st}$, otherwise considerable amounts of free labeled ligand would always be present and would interfere with measurements. $K_{d1}$ is the apparent $K_d$ for the labeled peptide used and has been experimentally determined as described in the previous paragraph. The FAM-labeled peptide was dissolved in dimethyl sulfoxide (DMSO) at 1 mM and diluted into experimental buffer. Readings were carried out with an Envision Multilabel Reader (PerkinElmer). Experiments were carried out in PBS (2.7 mM KCl, 137mM NaCl, 10 mM Na2HPO4 and 2 mM KH2PO4 (pH 7.4)) and 0.01% Tween 20 buffer. All titrations were carried out in triplicate. Curve-fitting was carried out using Prism 4.0 (GraphPad). To validate the fitting of a 1:1 binding model we carefully ensured that the anisotropy value at the beginning of the direct titrations between Mdm2 and the FAM-labeled peptide did not differ significantly from the anisotropy value observed for the free fluorescently labeled peptide. Negative control titrations of the ligands under investigation were also carried out with the fluorescently labeled peptide (in the absence of Mdm2) to ensure no interactions were occurring between the ligands and the FAM-labeled peptide. In addition, we ensured that the final baseline in the competitive titrations did not fall below the anisotropy value for the free FAM-labeled peptide, which would otherwise indicate an unintended interaction between the ligand and the FAM-labeled peptide to be displaced from the Mdm2 binding site.

#### Whole Cell Homogenate Stability

Peptides at a concentration of 1 μM were incubated at 37 °C with HCT116 whole cell homogenates prepared from 1 million lyzed cells/mL. The reaction was stopped at 0, 1, 2, and 4 hours and 22 hours with an organic solvent followed by centrifugation. The resulting supernatant was injected to LC/MS for the detection of tested peptide. The remaining percentage of each compound was normalized to the 0 hour amount and reported.

#### Plasma Stability

Peptide was incubated with human plasma at the concentration of 1 mM at 37 C for 1, 2, 3, and 4 hours. The incubation was stopped at indicated time points by addition of organic solvent followed by centrifugation. Parent compound in supernatant was analyzed by LC/MS. Percent remaining of peptide was calculated against the amount of compound at time 0.

#### p53 Beta-Lactamase Reporter Gene Cellular Functional Assay

HCT116 cells were stably transfected with a p53 responsive β-lactamase reporter and expanded in McCoy's 5A Medium with 10% fetal bovine serum (FBS), Blasticidin, and Penicillin/Streptomycin and then transferred to 1.5 ml freezing vials and stored under liquid nitrogen in growth media containing 5% DMSO. One day prior the assay, a vial of banked cells was recovered in a cell culture flask and incubated for 24 hours, followed by removal of cell growth media and replacement with Opti-MEM containing 2% FBS. The cells were then seeded into a 384-well plate at a density of 8000 cells per well. Peptides were then dispensed to each well using a liquid handler, ECHO 555, and incubated 16 h. The final working concentration of DMSO was 0.5%. β-lactamase activity was detected using the ToxBLAzer Dual Screen (Invitrogen), as per the manufacturer's instructions. Measurements were made using the Envision multiplate reader (Perkin–Elmer). Maximum p53 activity was defined as the amount of β-lactamase activity induced by 50 μM azide-ATSP-7041. This was determined as the highest amount of p53 activity induced by azide-ATSP-7041 from titrations on HCT116 cells.

#### Lactate Dehydrogenase (LDH) Release Assay

HCT116 cells were stably transfected with a p53 responsive β-lactamase reporter and expanded in McCoy's 5A Medium with 10% fetal bovine serum (FBS), Blasticidin, and Penicillin/Streptomycin and then transferred to 1.5 ml freezing vials and stored under liquid nitrogen in growth media containing 5% DMSO. One day prior the assay, a vial of banked cells was recovered in a cell culture flask and incubated for 24 hours, followed by removal of cell growth media and replacement with Opti-MEM containing 2% FBS. The cells were then seeded into a 384-well plate at a density of 8000 cells per well. Peptides were then dispensed to each well using a liquid handler, ECHO 555, and incubated 16 h. The final working concentration of DMSO was 0.5%. Lactate dehydrogenase release was detected using the CytoTox-ONE Homogenous Membrane Integrity Assay Kit (Promega), as per the manufacturer's instructions. Measurements were carried out using the Tecan plate reader. Maximum LDH release was defined as the amount of LDH released as induced by the lytic peptide (iDNA79) and used to normalize the results.

**Tetracycline Beta-Lactamase Reporter Gene Cellular Assay (Counterscreen)**

This assay was based on Jump-In™ T-REx™ CHO-K1 BLA cells containing a stably integrated β-lactamase under the control of an inducible cytomegalovirus (CMV) promoter. Cells were maintained in Dulbecco's Minimal Eagle Medium (DMEM) with 10% fetal bovine serum (FBS), Blasticidin, and Penicillin/Streptomycin and then transferred to 1.5 ml freezing vials and stored under liquid nitrogen in growth media containing 5% DMSO. One day prior the assay, a vial of banked cells was recovered in a cell culture flask and incubated for 24 hours, followed by removal of cell growth media and replacement with Opti-MEM containing 2% FBS. Cells were seeded into a 384-well plate at a density of 4000 cells per well. Peptides were then dispensed to each well using a liquid handler, ECHO 555 and incubated for 16 h. The final working concentration of DMSO was 0.5%. β-lactamase activity was detected using the ToxBLAzer Dual Screen (Invitrogen), as per the manufacturer's instructions. Measurements were carried out using the Envision multiplate reader (Perkin–Elmer). Counterscreen activity was defined as the amount of β-lactamase activity induced by tetracycline.

**Yeast 2-hybrid (Y2H) assay**

The Y2H assay was performed as previously described [14]. Briefly, yeast cultures that were grown overnight to saturation in SD-Leu-Trp medium were washed and diluted at $OD_{600}$ =0.2 in 200 μL SD-Leu-Trp-His-Ade medium containing the desired concentration of the peptide in duplicatein a 96- well microplate. The cultures were then placed in a 30 °C incubator with shaking for 3 days. Growth of the cultures was measured by recording the $OD_{600}$ at different time points using a Gen 5™(BIO-TEK Instrument, Vermont, USA) microplate reader. Average of the duplicate $OD_{600}$ readings was calculated and used in the analysis.

**Isothermal Titration Calorimetry (ITC)**

Overnight dialysis of protein and peptides were carried out in buffer containing 1× phosphate-buffered saline (PBS) pH 7.2, 3% DMSO, and 0.001% Tween-20. Approximately 100–200 µM of peptide was titrated into 20 µM of purified recombinant human Mdm2 protein (amino acids 1–125), over 40 injections of 1 µL each. Reverse ITC (200 µM of Mdm2 protein titrated into 20 µM of peptide) was carried out for peptides that are insoluble at high concentrations. All experiments were performed in duplicates using the MicroCal PEAQ-ITC Automated system. Data analysis was carried out using the MicroCal PEAQ-ITC Analysis Software.

**Circular Dichroism (CD)**

A total of 5 µL of the 10 mM stock peptide was mixed with 45 µL of 100% methanol, and dried for 2 h in the SpeedVac concentrator (Thermo Scientific). The dried peptide was reconstituted in a buffer (1 mM Hepes pH 7.4 and 5% methanol) to a concentration of 1 mM. The peptide sample was placed in a quartz cuvette with a path length of 0.2 cm. The peptide concentration was determined by the absorbance of the peptide at 280 nM. The CD spectrum was recorded from 300 to 190 nm using the Chirascan-plus qCD machine (Applied Photophysics, Surrey, UK), at 25 °C. All experiments were done in duplicates. The CD spectrum was converted to mean residue ellipticity, before deconvolution and estimation of the secondary structure components of the peptide using the CDNN software (distributed by Applied Photophysics).

**Surface Plasmon Resonance (SPR)**

SPR experiments were performed with Biacore T100 (GE Healthcare) at 25 °C. The site-specific mono-biotinylated Mdm2 was prepared by sortase-mediated ligation. The SPR buffer consisted of 50 mM Tris pH 7.4, 150 mM NaCl, 1 mM DTT, 0.05% Tween 20, and 3% DMSO. The CM5 chip was first conditioned with 100 mM HCl, followed by 0.1% SDS, 50 mM NaOH, and then water, all performed twice with 6 sec injection at a flow rate of 100 µL/min. With the flow rate set to 10 µL/min, streptavidin (S4762, Sigma-Aldrich) was immobilized on the conditioned chip through amine coupling, as described in the Biacore manual. Excess protein was removed by 30 s injection of the wash solution (50 mM NaOH + 1 M NaCl) at least eight times. The immobilized level was ~3000 RU. The biotinylated Mdm2 was captured by streptavidin, up to a level of ~400 RU. A flow cell consisting of only streptavidin was used as the reference surface. Using a flow rate of 30 µL/min, peptides dissolved in the SPR buffer were injected for 180 s. The dissociation was monitored for 300 s. For each peptide concentration, the peptide injection was followed by a similar injection of SPR buffer to allow the surface to be fully regenerated (though not completely for peptides with an extremely slow off-rate). After the run, responses from the target protein surface were transformed by: (i) correcting with the DMSO calibration curve, (ii) subtracting the responses obtained from the reference surface, and (iii) subtracting the responses of the buffer injections from those of peptide injections. The last step is known as double referencing, which corrects the systematic artefacts. The resulting responses were subjected to kinetic analysis by global fitting with a 1:1 binding model to obtain the *KD*, $k_a$ (M-1 s-1), and $k_d$ (s-1). Binding responses that did not have enough curvature during the association and dissociation were subjected to steady-state analysis, and the *KD* was obtained by fitting a plot of response at equilibrium against the concentration.

**HCT-116 Western blot analysis**

**Preparation of compound Stock and working Solutions**: 10 mM or 1 mM stock solutions of compounds were prepared in 100% DMSO. Each compound was then serially diluted in 100% DMSO and further diluted 10-fold into HPLC grade sterile water to prepare 10X working solutions in 10% DMSO/water of each compound. Depending on the required volume used in the relevant assay, compounds were added to yield final concentrations as indicated in the relevant figure with a residual DMSO concentration of 1% v/v.

HCT116 cells (Thermo Fisher Scientific) were cultured in DMEM cell media, which was supplemented with 10% foetal calf serum (FBS) and penicillin/streptomycin. All cell lines were maintained in a 37 °C humidified incubator with 5% $CO_2$ atmosphere. HCT116 cells were seeded into 96 well plates at a cell density of 60,000 cells per well and incubated overnight. Cells were also maintained in DMEM cell media with 10% fetal bovine serum (FBS) and penicillin/streptomycin. Cell media was then removed and replaced with cell media containing the various compounds/vehicle controls at the concentrations indicated in DMEM cell media with 2% FCS. After the stated incubation time (4 or 24 hours) cells were rinsed with PBS and then harvested in 100 µl of 1x NuPAGE LDS sample buffer supplied by Invitrogen (NP0008).

Samples were then sonicated, heated to 90 °C for 5 mins, sonicated twice for 10 s and centrifuged at 13, 000 rpm for 5 minutes. Protein concentrations were measured by BCA assay (Pierce). Samples were resolved on Tris-Glycine 4-20% gradient gels (BIORAD) according to the manufacturer's protocol. Western transfer was performed with an Immuno-blot PVDF membrane (Bio-Rad) using a Trans-Blot Turbo system (BIORAD). Western blot staining was then performed using antibodies against actin (AC-15, Sigma) as a loading control, p21 (118 mouse monoclonal), Mdm2 (2A9 mouse monoclonal) and p53 (DO-1 mouse monoclonal).

**Crystallization and data Collection**

Mdm2(6-125) was concentrated to 4.6 mg/mL and then incubated with thedouble stapled peptide at a 1:3 molar ratio of protein to peptide at 4°C overnight. The lyophilized double stapled peptide Mdm2-$^{d}$PMI-δ(1-5, 9-12) was first dissolved in DMSO to make a 100 mM stock solution before direct addition to the protein solution. The sample was clarified by centrifugation before crystallization trials at 16°C using the sitting drop vapour diffusion method. Crystals of Mdm2(6-125) in complex with $^{d}$PMI-δ(1-5, 9-12) were obtained by mixing the protein-peptide complex with the reservoir solution in a ratio of 1:1, with the reservoir solution containing 800 mM Sodium di-hydrogen phosphate, 800 mM di-Potassium hydrogen phosphate, 100 mM HEPES pH 7.5. Mdm2-$^{d}$PMI-δ(1-5, 9-12) complex crystals were frozen in an equivalent mother liquor solution containing 15% (v/v) glycerol and then flash frozen in liquid nitrogen. X-ray diffraction was collected at the Australian synchrotron (Aus.). See Table 3 for data collection statistics.

**Structure Solution and Refinement**

X-ray datasets were processed and scaled with the XDS[15] and CCP4 [16] packages. The structures were solved by molecular replacement with the program PHASER [17] using the human Mdm2 (6-125) structure from the PDB:4UMN (chain A) as a search model. The starting model was built and refined by iterative cycles of manual and automatic building with Coot [18] and restrained refinement with Refmac [19]. The d-amino acid peptide was manually buit into the electron density using COOT [18]. The geometric restraints for the non-natural amino acids constituting the hydrocarbon staples and the covalent bond linking their respective side chains together, to form the macrocyclic linkages constraining the stapled peptide, were defined and generated using JLigand [20]. The final model was validated using RAMPAGE

[21] and the MOLPROBITY [22] webserver. Structural overlays and analysis was performed using PYMOL[11]. See table 3 for data collection and refinement statistics. The Mdm2-$^{d}$PMI-δ(1-5, 9-12) complex structure has been deposited in the PDB with following accession code **XXXX.**
